# Tales of schizomid tails: patterns in schizomid flagellum shape from elliptical Fourier analysis

**DOI:** 10.1101/2021.10.05.463216

**Authors:** Robert J. Kallal, Gustavo Silva de Miranda, Erika L. Garcia, Hannah M. Wood

**Affiliations:** Department of Entomology, National Museum of Natural History, Smithsonian Institution, 10th & Constitution Ave, Washington, DC 20560; Department of Zoology, Denver Museum of Nature & Science, 2001 Colorado Blvd, Denver CO 80205; Department of Integrative Biology, University of Colorado Denver, 1201 Larimer St, Denver CO 80204

**Keywords:** arachnid, biogeography, Gaussian clustering, geometric morphometrics, outlines

## Abstract

The arachnid order Schizomida is a relatively understudied group of soil-dwelling predators found on all continents except Antarctica. While efforts to understand their biology are growing, there is still much to know about them. A curious aspect of their morphology is the male flagellum, a sexually dimorphic, tail-like structure which differs in shape across the order and functions in their courtship rituals. The flagellar shape is important for taxonomic classification, yet few efforts have been made to examine shape diversity across the group. Using elliptical Fourier analysis, a type of geometric morphometrics based on outline shape, we quantified shape differences across a combined nearly 550 outlines in the dorsal and lateral views, categorizing them based on genus, family, biogeographic realm, and habitat, with special emphasis on Caribbean and Cuban fauna. We tested for allometric relationships, differences in disparity based on locations and sizes in morphospace among these categories, and for clusters of shapes in morphospace. We found multiple differences in all categories despite apparent overlaps in morphospace, evolutionary allometry, and evidence for discrete clusters in some flagellum shapes. This study can serve as a foundation for further study on the evolution, diversification, and taxonomic utility of the male flagellum.

## Introduction

Schizomida is one of the less speciose orders of arachnids (scorpions, spiders, mites, and their relatives), with approximately 300 described species^1,2^ and 72 described genera (Supplement S1). They are small (2–12.5 mm), active hunters exhibiting sexual or asexual reproduction. Schizomids live in forests, caves, or synanthropic areas on all continents except Antarctica, but in general they are most diverse in warmer, tropical climates, with decreasing diversity as latitude increases. To date, the majority of species are described from Australia, Cuba, and Mexico^3,4^, which may be a reflection of the distribution of schizomid taxonomic expertise rather than a real biological effect. Schizomids are classified into two families - Protoschizomidae and Hubbardiidae - which together are the sister lineage of the larger, better known thelyphonids, colloquially called vinegaroons or whip scorpions. This relationship between schizomids and thelyphonids has been established morphologically^5,6^ and recently corroborated by high-throughput molecular analyses^7–11^. Relationships within Schizomida are recently being better understood, with phylogenetic analyses focusing on specific clades (e.g.^12^) or larger patterns^13^, but much remains to be done not only on their taxonomy, but on basic knowledge of their biology.

Among the traits shared between schizomids and thelyphonids is the presence of a flagellum, i.e., a short, tail-like, segmented structure. Female schizomids have an annulated flagellum, similar to that of thelyphonids in miniature, giving them the English colloquial name ‘short-tailed whip scorpion’ (Fig 1B). But there is sexual dimorphism in this structure, with the male flagellum being a non-annulated, stouter, more complex structure that is used during courtship (Fig. 1A)^14,15^. Despite its importance to species diagnoses^16,17^, only recently attempts have been made at homologizing aspects of the flagellum, namely annuli, setae, lobes, and depressions, in the hope that this knowledge will clarify evolutionary relationships. Those studies, however, focus on more specific lineages or solely on females^2,18^, leaving out the shape and size of the male flagella, which varies widely across schizomids. They differ in length by nearly an order of magnitude, ranging from 0.2 mm in *Adisomus duckei* Cokendolpher and Reddell, 2000 and *Rowlandus littoralis* Teruel, 2003 (∼7% total body length) to 1.68 mm in *Piaroa villarreali* Armas and Delgado-Santa, 2012 (∼30% of 5.56 mm total body length). The male flagellar shape is phenotypically diverse and characterizing them for quantitative analysis has proven difficult. Linear morphometric ratios^19^ and discrete states^20^ have been used to code them for cladistic analyses, although, doing so reduces the available shape diversity. Thus, the male flagellum is critical for classification and species identification, and is likely important for sexual selection and reproduction, yet, there has been little research done that examines the shape diversity across schizomids^4,17^.

**Figure 1.**
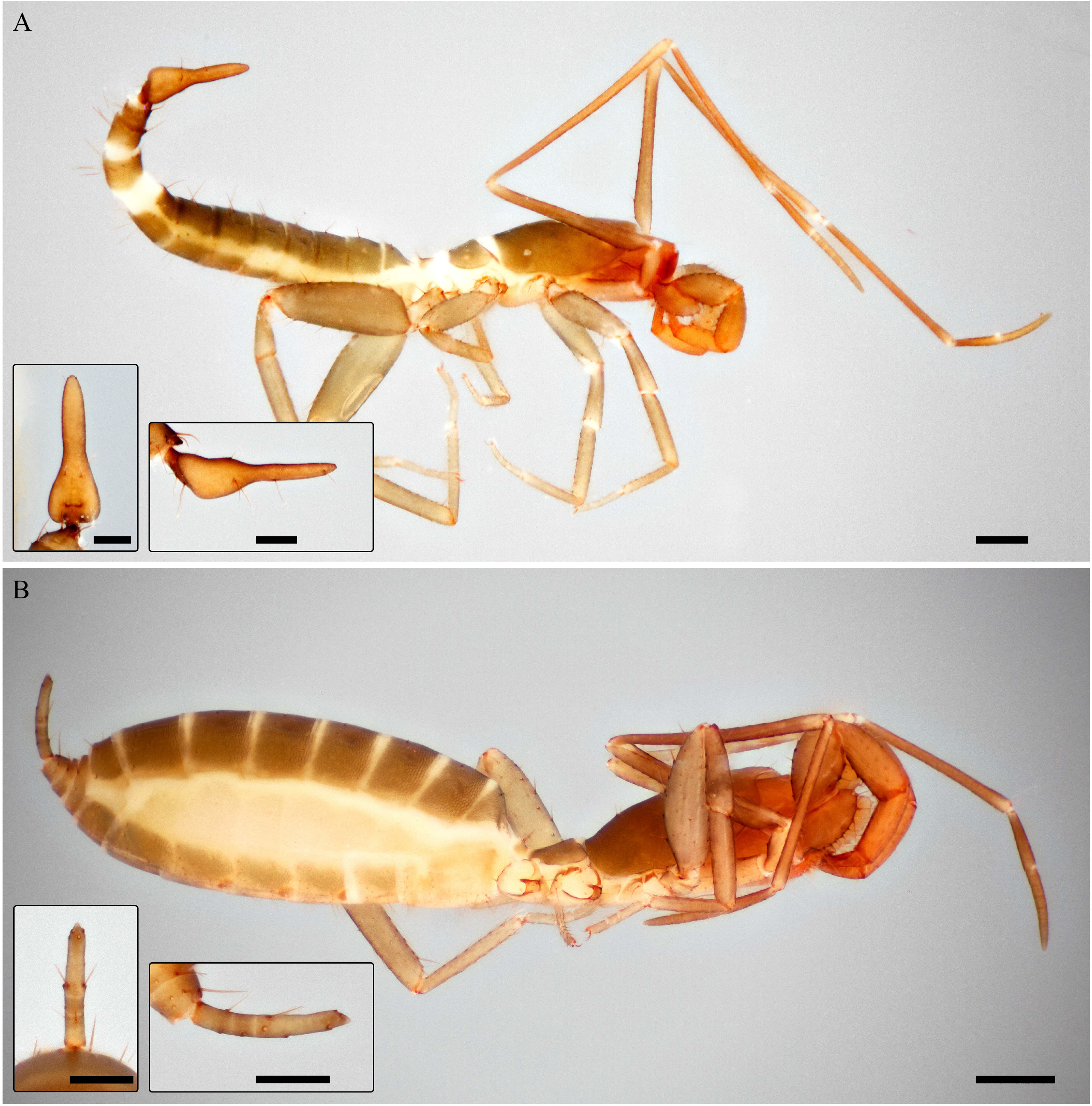
Habitus and flagellum images of *Hubbardia pentapeltis* Cook, 1899. A. Male (USNMENT 1538005), lateral; inset left, flagellum in dorsal view; inset right, flagellum in lateral view. B. Female (USNMENT 1458794), lateral; inset left, flagellum in dorsal view; inset right, flagellum in lateral view.

One issue is that landmark-based morphometric analysis in flagellum shapes is limited. This is due to the high variability in shapes and the absence of certain structures from some flagella, which prevents assigning landmarks based on a homology criterion. A solution is elliptical Fourier analysis, which differs from landmark-based geometric morphometrics by quantifying 2- dimensional outlines using sine and cosine functions, thus allowing for study of structures that lack homologous landmarks^21,22^. The accumulation of those functions, called harmonics, captures a shape in increasing detail. From the harmonics, Fourier coefficients are generated, which may be subjected to downstream multivariate analyses. This type of morphometric analysis has been conducted on other arachnid groups where shape variability has precluded the use of landmark-based methods: the lateral profile of mygalomorph spiders’ carapaces^23^, the dorsal scutum of podoctid opiliones^24^, the raptorial limbs of whip spiders^25^, and solifuge chelicerae (Garcia & Cushing, in prep). To our knowledge, no such analysis has been conducted on schizomids.

Here, we aim to characterize the major axes of variance in male schizomid flagellum shape using elliptical Fourier morphometrics, sampling across schizomids for more than 80% of the known species diversity. Our work capitalizes on the evolutionary information extractable from the primary taxonomic literature (e.g.,^26^), with the vast majority of outlines obtained from published descriptions. In order to understand how shape changes with respect to size we tested for an allometric relationship between flagellum shape and size. In order to identify clades, areas, or habitats with high flagellum shape diversity, we tested for differences in disparity, or phenotypic diversity, based on genus, biogeographic realm, and microhabitat, with disparity analyzed by area and location metrics. Then we focused on the Caribbean schizomofauna - in particular, Cuba - as a subset to look for correlations of island size with disparity and diversity. Given the flagellum is important to courtship, one expects they may be more variable on islands with more species to avoid hybridization, with higher variability measured as a disparity with a higher area occupied in morphospace. Finally, we use Gaussian mixture modeling to observe if there are reliable clusters of flagellum shapes. In this study we characterize for the first time the shape diversity present in a schizomid sexual structure, and highlight important patterns in schizomid morphological diversity.

## Methods and material

### Outline acquisition and classifiers

Schizomid species descriptions typically include a dorsal, lateral, and, less often, ventral view of the male flagellum (e.g., Fig1A, inset). A total of 547 images - 283 dorsal and 264 lateral - were selected (e.g., Supplements 2-6). This includes 64 genera with dorsal images, which comprises all genera with an illustrated male, sampling across ∼90% of described schizomid diversity at the genus level and ∼80% of described schizomid species diversity. These images were converted to silhouettes in Adobe Photoshop and GIMP. Illustrations rotated from a standard dorsal or lateral views were omitted from analyses as preliminary analyses interpreted resulting asymmetry as real differences in shape rather than artifacts. In the case of dorsal images, the left side was mirrored to remove asymmetry artifacts in illustrations. Reported flagellum lengths were used for scaling.

Data for classifying outlines into subsets were assembled into a comma-delimited files (Supplements 7-10). These include genus and family level taxonomic classification, habitat type (epigean or hypogean), country of distribution, and biogeographic realm. In the latter case, taxa were classified as Australasian, Afrotropical, Indomalayan, Palearctic, Nearctic, Neotropical, or from the Mexican Transition Zone. The latter is the interface between the Nearctic and Neotropical realms found in Mexico, an important biogeographic area where many schizomids have been collected^27,28^. These data are summarized in Table 1.

**Table 1.**
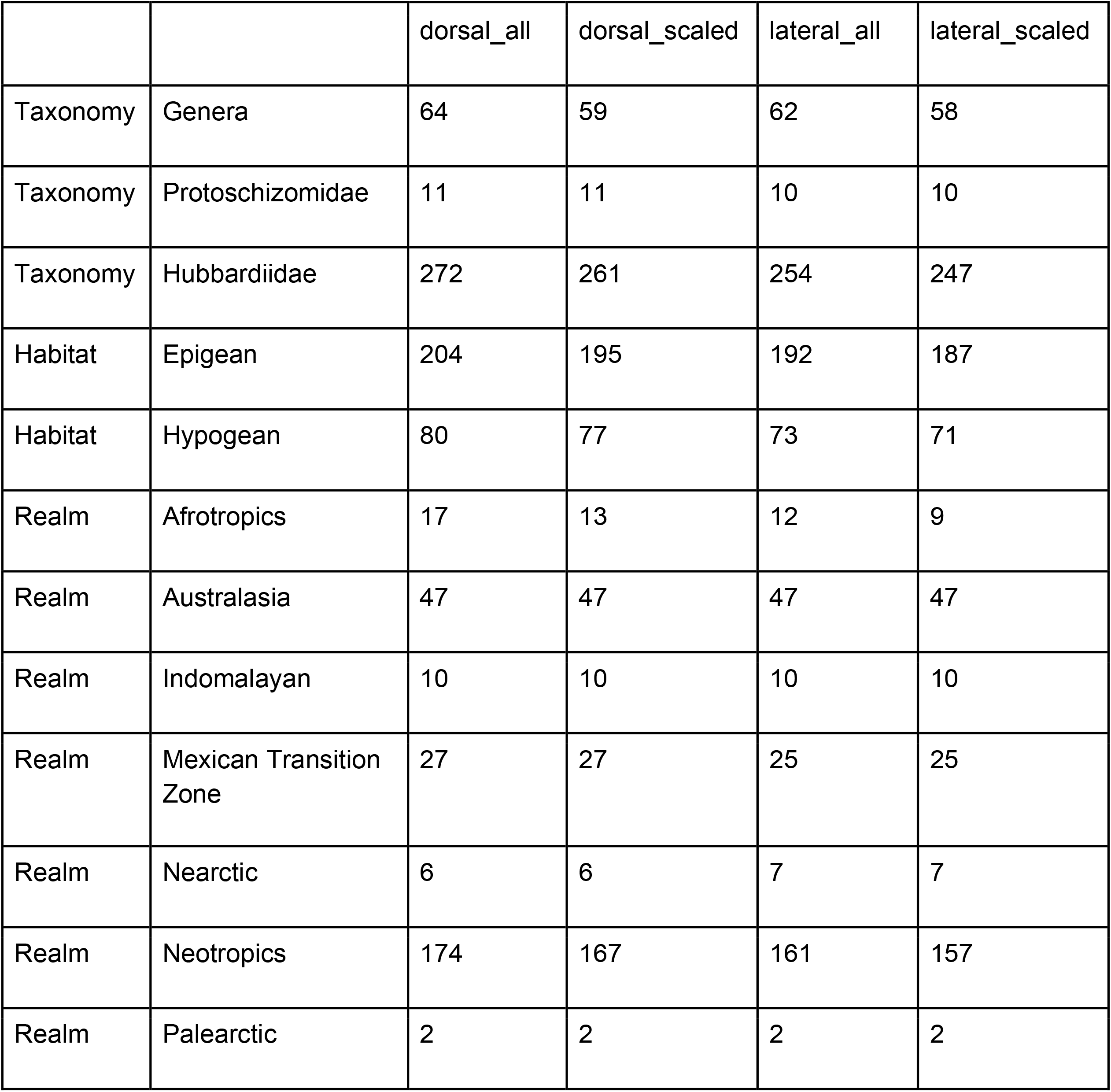
Number of schizomid flagellum outlines in each global dataset, categorized by taxonomy, habitat, and biogeographic realm.

### Dataset preparation and elliptical Fourier analyses

The elliptical Fourier analysis pipeline uses the package *Momocs*^29^ in R^30^. Outlines were smoothed, scaled to remove size, centered, and the starting point was slid to the upper median position, forming the *dorsal_all* dataset, of which three other datasets were derived. A reduced dataset of 272 dorsal outlines with associated flagellum length data was analyzed as above but using the flagellum length to scale the outline sizes to their actual sizes, rather than minimized as in typical morphometric analyses; this is the *dorsal_scaled* dataset. Two reduced datasets based on geography, with only Caribbean and Cuban species, were analyzed and called *dorsal_caribbean* and *dorsal_cuba*, respectively. A sister set of analyses of the lateral shapes was conducted on lateral outlines (*lateral_all*), 257 scaled lateral outlines (*lateral_scaled*), outlines of Caribbean taxa (*lateral_caribbean*), and outlines of Cuban taxa (*lateral_cuba*). Analyses were conducted on dorsal datasets unless otherwise noted. Harmonic power was calibrated to capture 99.9% of variation and the elliptical Fourier analysis was performed using the recommended number of harmonics using the *Momocs* function ‘calibrate_harmonicpower_efourier’.

### Multivariate analyses

A principal components analysis (PCA) was conducted on the resulting Fourier coefficients. PC axes that explained more than 1% of the variation were retained. Shape variation along the axes were visualized using the ‘PCA’ and ‘PCcontrib’ functions in *Momocs*. Morphospace plots were generated using the R package *borealis*^31^.

To test for an allometric relationship between flagellum size and shape, we generated centroid sizes on the *scaled* datasets and PCs explaining more than 1% of variance in the *all* datasets. A linear model to test if the log transformed centroid size is a predictor of shape was conducted in the R package *RRPP*^32,33^. Disparity analyses were conducted using the R package *dispRity*^34^ to compare differences in phenotypic variation among taxonomic units, areas and habitats, following^35^. An area metric, the sum of variances, was used to measure the hypervolume area and a position metric, the distance of observations from the centroid, was used to measure the location of a hypervolume in morphospace^36,37^. PCA shape scores were bootstrapped with 100 replicates and rarefied to the size of the smallest subset; subsets were removed when represented by less than three entries. This equalizes the sampling but may result in subsampling of larger groups that does not fully capture their disparity. A Wilcoxon signed rank test was used for the comparison of disparity values, with a Bonferroni correction in the case of multiple comparisons using the ‘test.disprity’ function in *dispRity*. Pearson correlations were conducted using the base R functions^30^, comparing both species richness and disparity of genera worldwide as well as on a subset Caribbean islands with schizomids sampled here - Cuba, Hispaniola, Puerto Rico, Jamaica, Navassa, and Trinidad and Tobago. Island land area was also tested for correlations with the latter.

To test for the taxonomic and cladistic utility of discrete states of the male flagellum, we used Gaussian mixture modeling in the R package *mclust*^38^. Gaussian finite mixture models, fitted by expectation-maximization (EM) algorithm, are evaluated via Bayesian information criterion (BIC) to determine the best-fit clustering model. The first three principal components, which explain more than 95% of variation, were used as data input. Individual outlines of uncertain cluster membership were removed from clusters at a 0.05 and 0.25 threshold, due to uncertainty values ranging from 0 (highly certain) to 1 (highly uncertain). Mean shapes of resulting clusters were determined by the function MSHAPES in Momocs, and a Pearson correlation was conducted to examine the relationship between number of individuals in a genus versus the number of clusters they occupy.

All scripts can be found at https://github.com/bobkallal/schizomida.

## Results

### PCA loadings and axes of variation

A total of 17 harmonics explained 99.9% of the variation in the *dorsal_all* dataset, resulting in 68 PC axes. The first six individually explained more than 1% of variation each, for a total of 97.3%. The first PC’s extremes are slender *Colombiazomus truncatus* Armas and Delgado-Santa, 2012 and wide *Kenyazomus pekkai* Armas, 2014, and explain 71.9% of variation. The second PC ranges from the spade-shaped *Bamazomus subsolanus* Harvey, 2001 with a long pedicel to *Schizomus procerus* (Hansen in Hansen and Sörensen, 1905), which is widest anteriorly and a short pedicel; this axis explains 13.4% of variation. The third PC, which explains 5.7% of variation, has a broad pedicel with posterior-directed lateral lobes in *Protoschizomus sprousei* Cokendolpher and Reddell, 1992 to *B. vespertinus* Harvey, 2001, with anterior-directed lateral lobes. The fourth PC’s extremes are broad and square, as in *Surazomus manaus* Cokendolpher and Reddell, 2000, to the trident-shape of *Cokendolpherius ramosi* Armas, 2002; it explains 3.6% of variation. The *dorsal_scaled* dataset’s first PC broadly corresponds to length, ranging from *Cubazomus montanus* Teruel, 2004 to *Hansenochrus urbanii* Villarreal and Teruel, 2006 and accounting for 79.9% of variance; PCs 2–4 correspond to PCs1–3 in the *dorsal_all* analysis, and explain 15.1%, 2.4%, and 1.0% variance, respectively (Supplement 11).

In the *lateral_all* dataset, a total of 16 harmonics explained 99.9% of the variation, thus resulting in 64 PCs. The first two PCs explain 50.6% and 23.5% of variation, respectively (Figs. 2B, 3B, 3D). As in the dorsal analysis, the first PC explains relative slenderness, ranging from *Piaroa virichaj* Villarreal, Giupponi and Tourinho, 2008 to *Rowlandius ramosi*. The inflection of the flagellum is summarized by PC2, ranging from the downward *Troglocubazomus inexpectatus* Teruel and Rodríguez-Cabrera, 2019 to the upward *Schizomus hanseni* Mello-Leitão, 1931. The third PC explains if the flagellum is more convex dorsally or ventrally, with the extremes being *Notozomus majesticus* Harvey, 2000 and *Attenuizomus mainae* (Harvey, 1992); it includes 6.3% of variation. The last PC detailed here, PC4, summarized whether the pedicel or distal part of the bulb is thinner, ranging from *Bamazomus vespertinus* to *Draculoides mckechnieorum* Abrams & Harvey, 2020 and includes 5.2% of variation. As in the dorsal dataset, the first PC of *lateral_scaled* corresponds to size, explaining 83.8% of variance, ranging from the long *Hansenochrus urbanii* to the small *Adisomus duckei*, with variance explained by PC2 corresponding to PC1 in the *lateral_all* dataset, and so forth, with PCs 2-4 explaining 8.2%, 3.6%, and 1.2%, respectively (Supplement 12)

**Figure 2.**
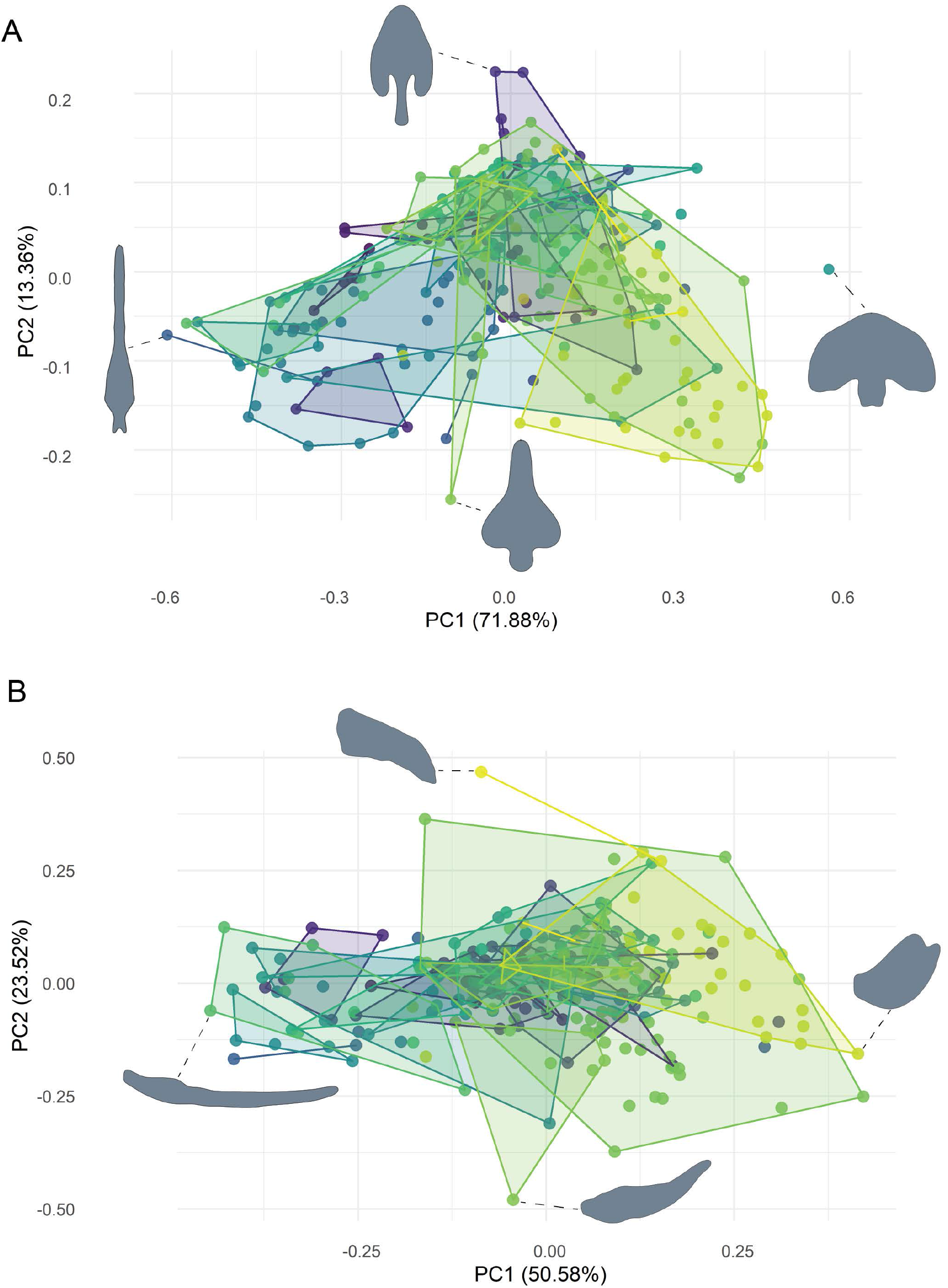
Morphospace plots based on male schizomid flagella, categorized by genus. A. Dorsal view. PC1 extremes are *Colombiazomus truncatus* Armas and Delgado-Santa, 2012 (negative) and *Kenyazomus pekkai* Armas 2014 (positive). PC2 extremes are *Schizomus procerus* (Hansen in Hansen and Sörensen, 1905) (negative) and *Bamazomus subsolanus* Harvey, 2001 (positive). B. Lateral view. PC1 extremes are *Piaroa virichaj* Villarreal, Giupponi and Tourinho, 2008 (negative) and *Rowlandius ramosi* Armas, 2002 (positive). PC2 extremes are *Schizomus hanseni* Mello-Leitão, 1931 (negative) and *Troglocubazomus inexpectatus* Teruel and Rodríguez-Cabrera, 2019 (positive).

**Figure 3.**
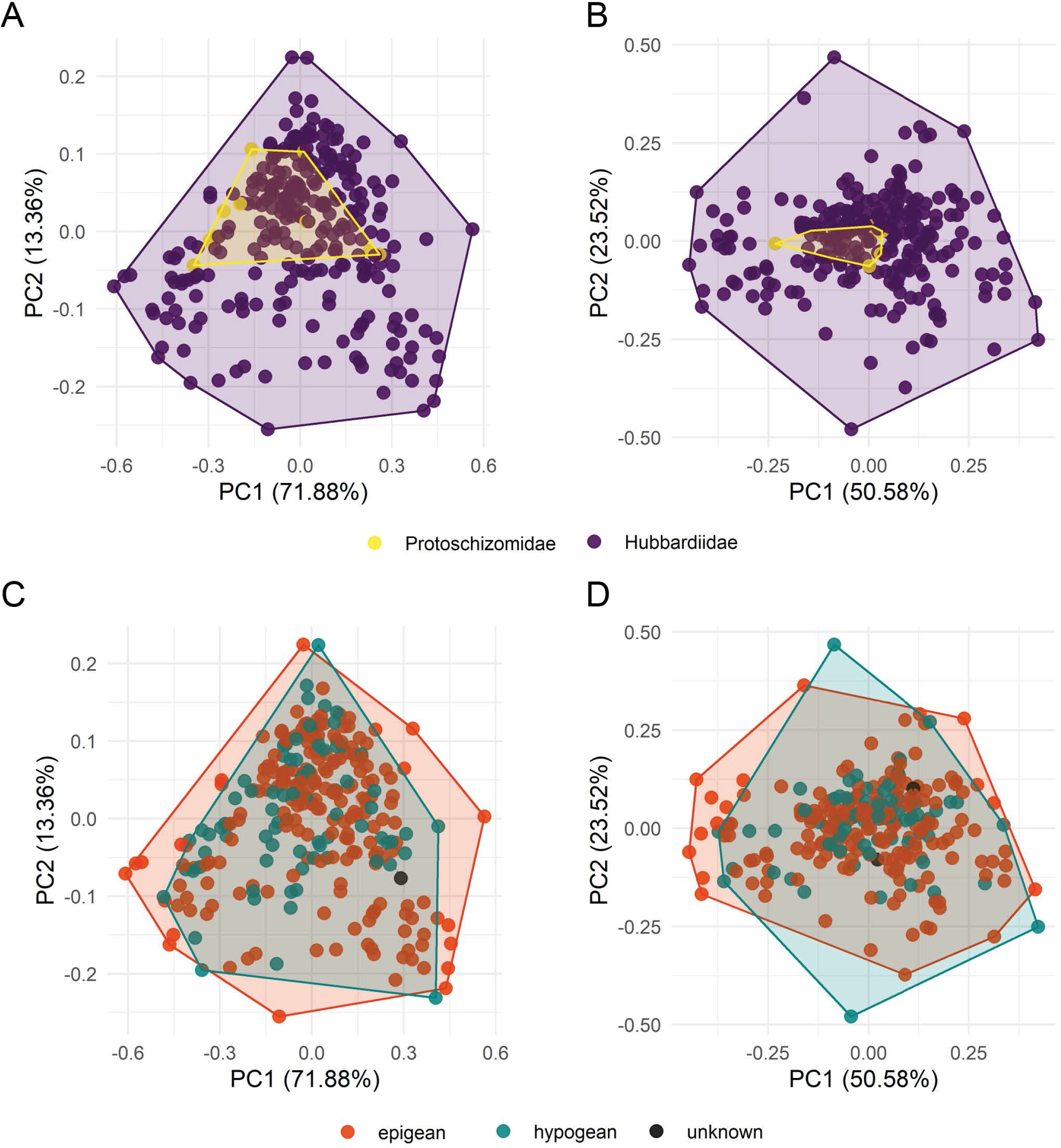
Morphospace plots based on male schizomid flagella. A. Categorized by family, dorsal view. B. Categorized by family, lateral view. Legend: Protoschizomidae, yellow; Hubbardiidae, violet. C. Categorized by habitat, dorsal view. D. Categorized by habitat, lateral view. Legend: epigean, red; hypogean: teal; unknown, black.

### Allometry and disparity

The linear model testing for allometry of the dorsal dataset resulted in R^2^ = 0.0759 and P = 0.001, and R^2^ = 0.0830 and P = 0.001 in the lateral dataset. The two families were not significantly different in the size of the area of morphospace occupied in the dorsal view (P = 0.218) but did significantly differ in the lateral view (P < 0.001). The location of the families’ centroids did not differ significantly in the dorsal view (P = 0.643) but did differ in the lateral view (P < 0.001). The area occupied by the two habitat types, epigean and hypogean, differed in area dorsally (P = 0.001) and laterally (P < 0.001) while their locations in morphospace did not differ from the dorsal aspect (P = 0.235) but did in the lateral aspect (P < 0.001). Most biogeographic realms differed significantly (P < 0.001) from each other in both area and location of morphospace occupied (Fig. 4). The test statistics for the Wilcoxon signed rank tests based on differences in sum of variances by family, biogeographic realm, and habitats are located in Supplement 13.

**Figure 4.**
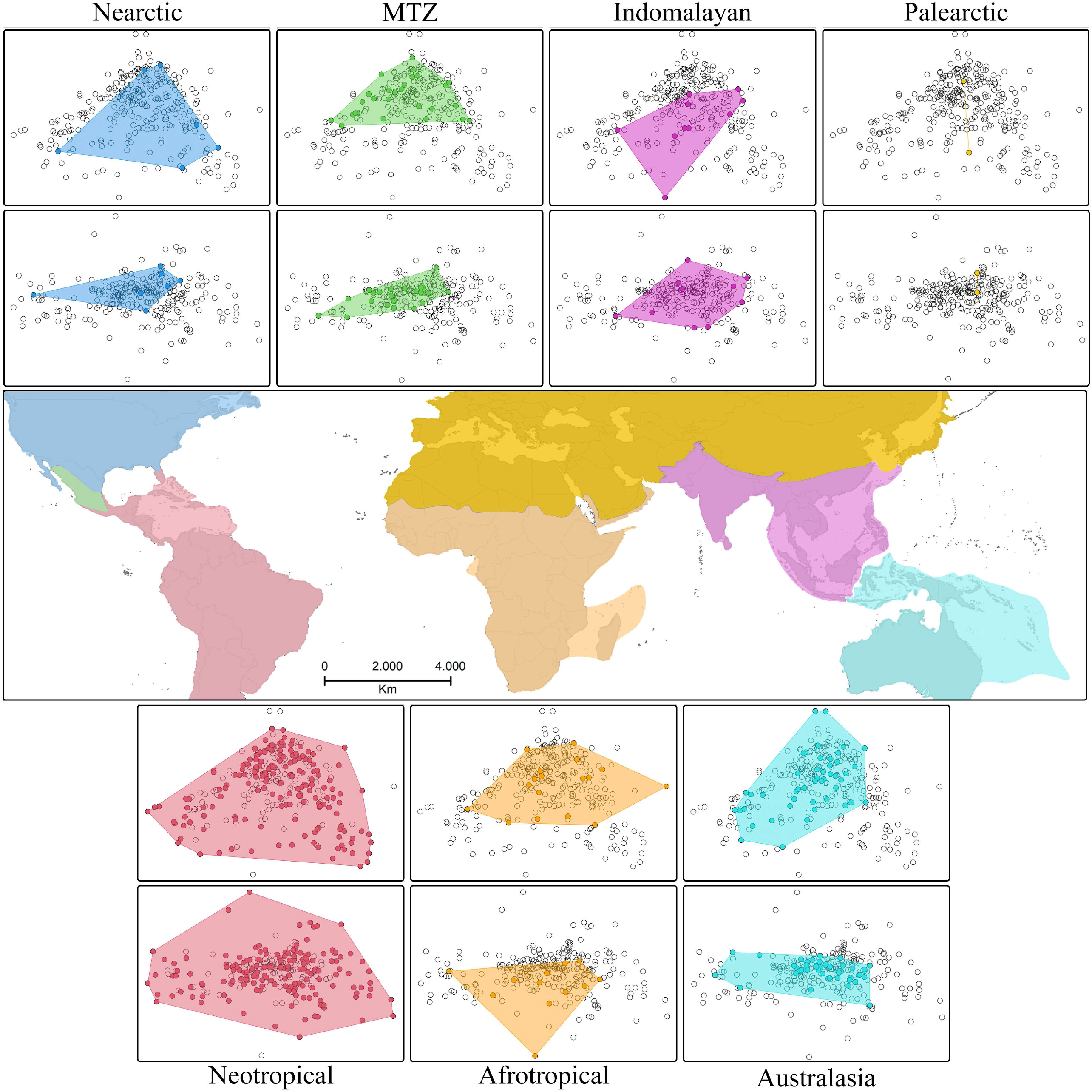
Morphospace plots in biogeographic realms based on male schizomid flagella. A. Nearctic. B. Mexican Transition Zone (MTZ). C. Indomalayan. D. Palearctic. E. Map shows locations of biogeographic zones. F. Neotropics. G. Afrotropics. H. Australasia.

Pearson’s correlation test for a relationship between island area and richness (based on *dorsal*_*all* as it has the best representation) resulted in R = 0.87 and P = 0.52. The test for correlation between island area and dorsal disparity resulted in R = −0.52 and P = 0.37 while the same for lateral disparity resulted in R = −0.38 and P = 0.52. We found the disparity (both area and location) found on most islands were significantly different from each other (P < 0.001) despite the apparent overlap in morphospace in both area and location occupied (Fig. 5). In the dorsal view, the areas for Puerto Rico and Jamaica were the same size, and the locations of Hispaniola, Jamaica, and Trinidad and Tobago were the same. In the lateral view, the areas of disparity of all islands differed and the only locations shared were between Jamaica and Trinidad and Tobago. No significant correlation was determined between dorsal disparity and individuals per genus across all genera (R = 0.096, P = 0.64).

**Figure 5.**
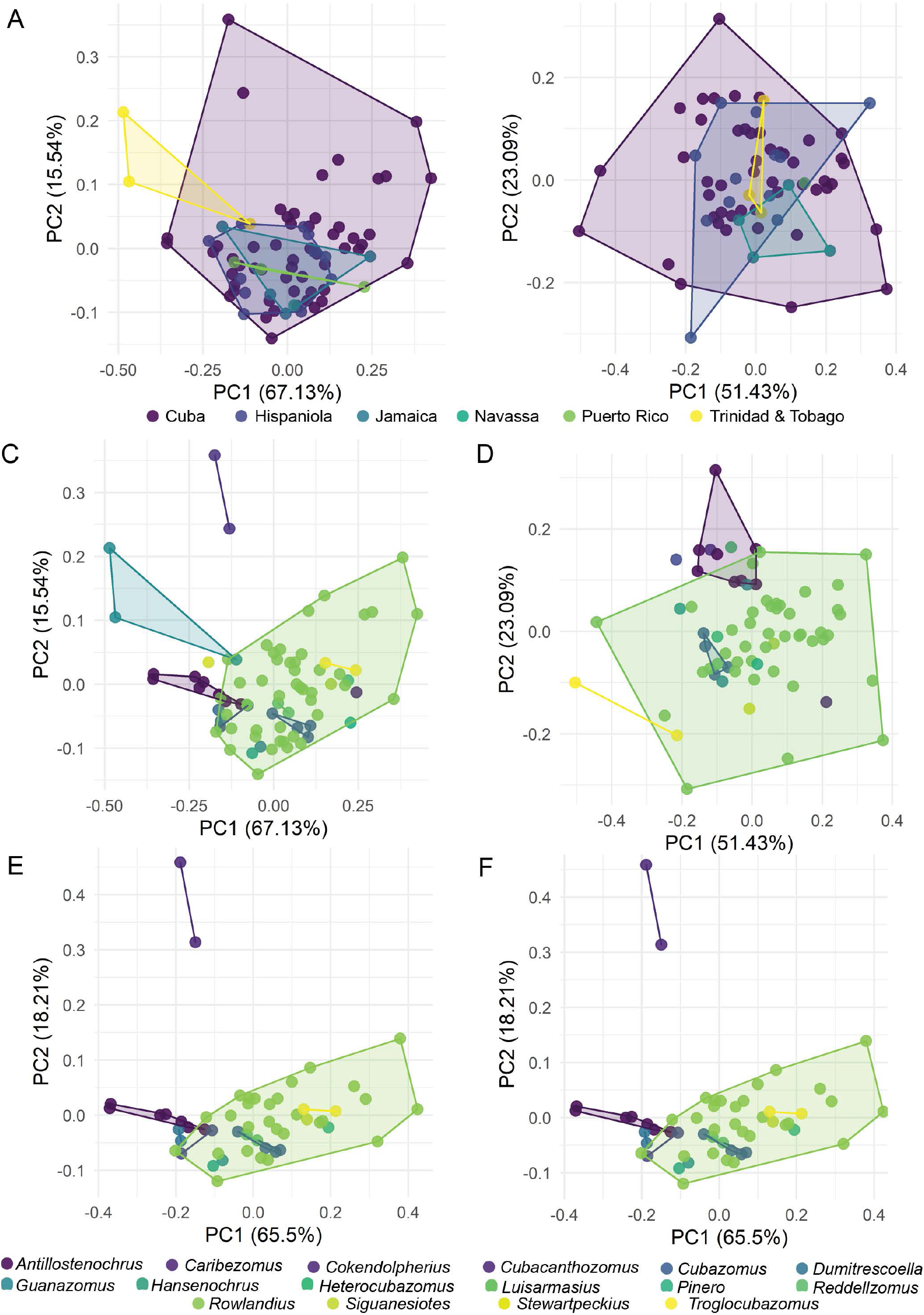
Morphospace plots based on male schizomid flagella of Caribbean and Cuban species. A. Dorsal, by island. B. Lateral, by island. C. Dorsal, by genus, from Caribbean islands. D. Lateral, by genus, from Caribbean islands. C. Dorsal, by genus, from Cuba. D. Lateral, by genus, from Cuba.

### Clustering

The best-fit model from the Gaussian mixture modeling was the ellipsoidal, equal volume, shape, and orientation (EEE) model with a log-likelihood of 932.118 and BIC of 1632.774. Nine clusters were supported, with a range of two to 102 members per cluster. Clusters were reduced by uncertainty levels of 0.25 and 0.05, where individuals with associated uncertainty values higher than the aforementioned thresholds were removed from the clusters. Using the 0.25 threshold, 54 outlines were of uncertain affinity (Fig. 6), and using the 0.05 threshold, 157 outlines could not be reliably clustered. Cluster one (black) includes 16-29% of flagella, more than any other cluster besides those uncertain flagella in the 0.05 threshold, with a typical subrhomboid shape with a pedicel about ¼-⅓ the width of the bulb and about as long. The second cluster (teal) is one of two with three distal lobes, with the median being the longest, and a relatively long pedicel. Cluster three (green) is spade-shaped, with recurved lateral lobes and elongated pedicel. Cluster four (brown) is lanceolate, with an elongate, tapering bulb and typical pedicel. Cluster five (cyan) is round with a very short pedicel. Cluster six (magenta) is somewhat rectangular, with little definition between bulb and pedicel. Cluster seven (yellow) is also trilobed, with three subequal lobes and virtually absent pedicel forming a badge-like shape. Cluster eight (red) has a rounded bulb with a vague point and typical pedicel. Cluster nine (blue) is triangular with a short pedicel. A significant correlation was found between number of individuals per genus and number of non-zero clusters occupied (R = 0.6, P < 0.001). Results for the stricter 0.05 threshold can be found in Supplement 13.

**Figure 6.**
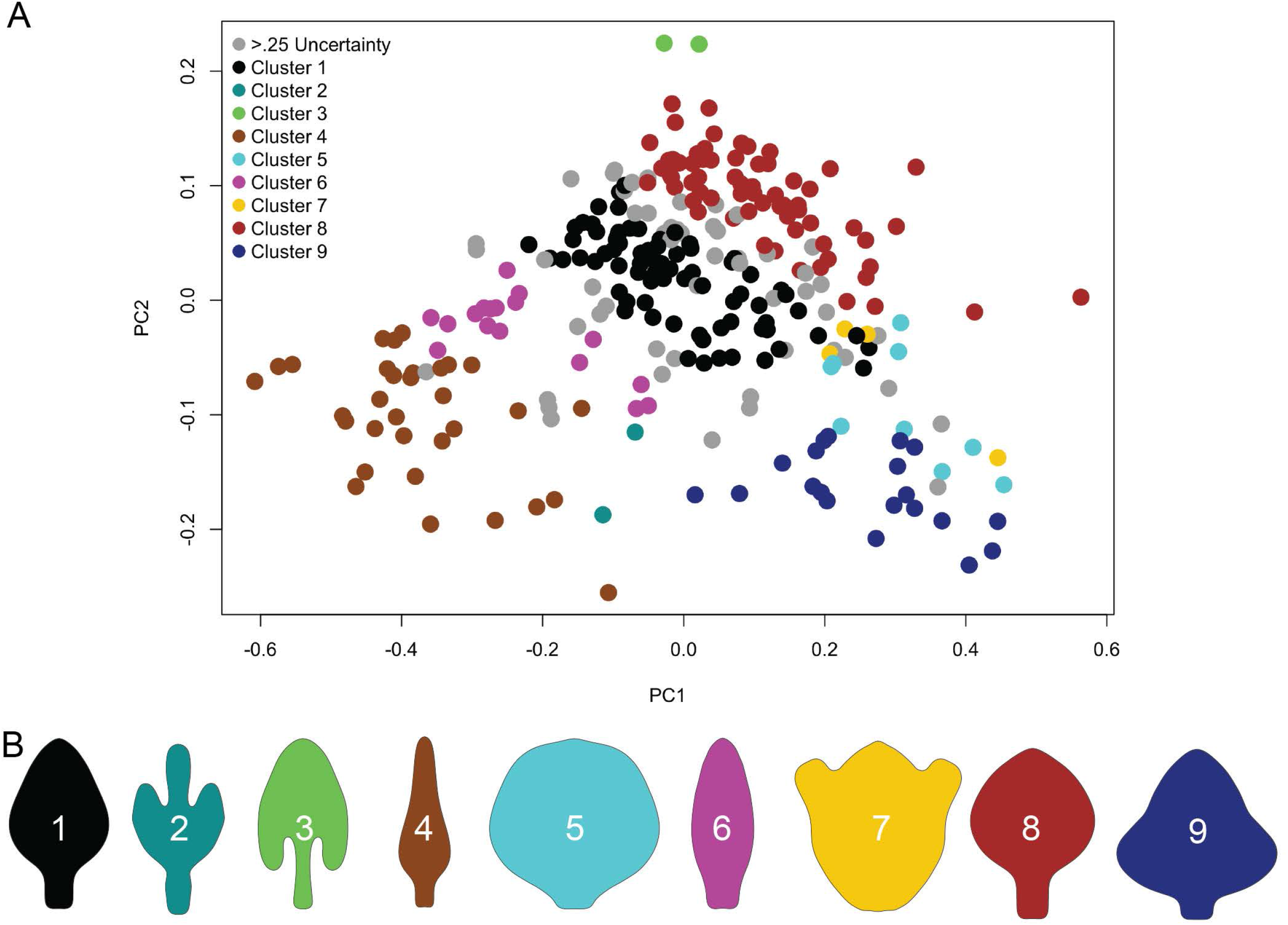
Gaussian clustering of male schizomid flagella in the dorsal view (*dorsal_all*). A. Morphospace plot with PC1 and PC2, and clusters demarcated by color. B. Mean shapes for respective clusters.

## Discussion

### Morphospace occupation and disparity

Morphospace plots showed broad overlap in sampled genera, which was unsurprising given their number and the continuous nature of their shapes. Despite the overlaps in morphospace among families, biogeographic realms, and habitats, we were still able to detect differences in area and location of morphospace of these groups in many cases. Area differences represent broad morphological diversity, and location differences imply a likelihood to have different shape space; the two disparity metrics are unlinked.

Differences in the area of morphospace occupied in the various biogeographic realms were detected, but the overlaps in morphospace between them led to fewer differences using the locations metric. This means the amount of variation differed but the shapes did not vary much between them, thus suggesting morphologies are not trending in different directions in different biogeographic realms. Differences in sample sizes in some areas, whether artefactual from differing sampling efforts or representing true differences in diversity also colors results, leading to the appearance of high disparity in low diversity realms (e.g., the Nearctic seems to have high disparity). Besides the Nearctic outlier, disparity was highest in the Neotropics, followed by the Afrotropics, then comparable disparity in the Australasian, Indomalayan, and Mexican Transition Zone faunae. This replicates schizomid distribution – where the bulk of species diversity is in the warm tropics, with less species diversity as you go away from the equator. The lateral disparity metrics were similar with the exception of a lower disparity for the fauna of the Mexican Transition Zone. When viewed dorsally, epigean schizomid flagella occupied a larger area of morphospace than hypogean ones, but their location in morphospace overlapped. From the lateral view, schizomids on and below the surface occupied different areas and locations of morphospace but they generally overlapped. In other words, we observed more shape difference dorsally in epigean species compared to hypogean species, but the types of shapes exhibited were similar. In contrast, the lateral outlines had different areas and different shapes, suggesting possible different pressures on courtship and mate recognition below the surface. The most notable place lacking difference was in the size and location of morphospace occupied by protoschizomids and hubbardiids in the dorsal view as compared to the lateral view, in which there was a difference. Protoschizomidae has much less species diversity than Hubbardiidae; only 11 of 283 dorsal outlines sampled here are protoschizomids, which approximately reflects the described species diversity. Despite the apparent size difference in areas occupied by protoschizomids and hubbardiids when comparing the first two PCs that explain the highest amount of variation, no area difference was statistically determined, principally due to *Protoschizomus sprousei* Cokendolpher and Reddell, 1992. Most representatives of *Protoschizomus* have a flagellum that is broader posteriorly, the bulb wider than the pedicel but with a smooth transition between them. Members of the other protoschizomid genus, *Agastoschizomus*, have similar flagella, albeit with more rectangular shapes. The flagellum of *P. sprousei* differs considerably from those, with a pair of lateral lobes. This shape extends the morphospace occupied by protoschizomids greatly, resulting in a comparable volume of morphospace occupied by the two families.

Despite the variability found in the male flagellum, the same cannot be said of its counterpart in the female chelicerae, which show no obvious variation. The chelicerae are the grasping structures of the mouthparts and during the mating march, she clasps his flagellum with her chelicerae as he searches for a place to deposit his spermatophore^14,15^. Curiously, the cheliceral shows little, much less commensurate, change, compared to the remarkable differences observed in the male spiders’ pedipalps (an intromittent organ) and the females’ external and internal genitalia. That is, no obvious arms race is apparent in external appearances. A reason for this may be the necessity of conservation of the chelicerae’s shape for other tasks such as feeding, which somehow does not preclude it from interlocking with the myriad shapes of male flagella, although presumably specialized setae, spines or grooves could have evolved that do not interfere with cheliceral function. Regardless, given this shape conservatism in the chelicerae, it is likely other mechanisms are driving flagellum shape differences in males.

### Allometry

We found signals of evolutionary allometry, with shape differences correlating with size in both the dorsal and lateral views. Hubbardiid species with more slender flagella (e.g., *Piaroa virichaj, Hubbardia pentapeltis* Cook, 1899, *Hansenochrus urbanii*) and the protoschizomid genus *Agastoshizomus*, members of which have largely rectangular flagella when viewed dorsally, have larger flagella. Flagella with a more typical rhomboid shape or with complex lobes tended to be smaller. A similar pattern is observed from the lateral view, with larger species being the slender hubbardiids and rectangular (but with a ventral process) *Agastoschizomus*. In both cases, these taxa seem to be driving the allometric relationship, with most shapes being smaller in size. This implies the relatively slim, elongated flagella may more easily get larger and more complex shapes apparently cannot.

### Schizomids in the Caribbean

In the more limited analysis of Caribbean taxa, we found mostly different morphospace areas per island but more overlap (Fig. 4A, B). That is, as measured here, the areas of morphospace occupied by the various islands differed, with some very high, like Cuba (51 species), and others unexpectedly low, as in the comparably sized Hispaniola (12 species). Meanwhile, most of the convex hulls overlapped, due in large part to the very large area containing the Cuban taxa in comparison to that of the other islands. In contrast, the area and location of the flagellum shapes of the various islands in morphospace did differ in the lateral view. However, as with the dorsal view, the diversity of the Cuba schizomofauna overlaps with the others, so despite different medians of centroids of their convex hulls, they all fall within Cuba’s convex hull. As it appears, the scope of shapes of male flagella in Cuba exceeds that of other Caribbean islands. The pattern of described diversity - with more on larger islands than smaller - is consistent with the theory of island biogeography^39^ and other terrestrial arthropods^40^. While we did not find a significant relationship between island size and described diversity, it is possible Trinidad and Tobago is more diverse due to its proximity to mainland South America and/or there is undercollection in Hispaniola. We also found no relationship between disparity and island area, which likely represents a difference in sample size rather than differences in morphospace volumes, similar to biogeographic realm comparisons (T. Guillerme, pers. comm.).

A major point of consideration when comparing Cuba’s schizomid diversity to other Caribbean islands is the relative preponderance of schizomid specialists. Teruel and de Armas are responsible for virtually all of the described Cuba taxa and the majority of the described taxa from the Dominican Republic. The relative dearth of described species from Haiti, Jamaica, Puerto Rico, and Trinidad and Tobago may therefore represent an artifact of taxonomic effort in those areas rather than reduced diversity in comparison to Cuba.

Because of this taxonomic effort, we can consider Cuba to represent a more complete sample of true morphological diversity in an area. We found the morphospace occupied by *Rowlandius* dwarfed that of the other genera, due at least in part to its increased species diversity compared to the other genera, some of which are monotypic (Fig. 5C-F). The three genera with more than three specimens - *Antillostenochrus, Cubazomus*, and *Rowlandius* - could be distinguished based on area and location in morphospace in both the lateral and dorsal views. This suggests that taxa in a taxon rich locale like Cuba may be distinguished based on shape, which may be a boon to parataxonomists seeking to catalogue the diversity found there. A well sampled, time-calibrated phylogeny would be key to understanding possibilities of convergent morphologies and cryptic species (or genera) as well as colonization timings.

### Clustering

The clustering algorithm suggested nine clusters for male schizomid flagella (Fig. 6). Clusters two, three, four, five, seven, and nine are essentially unchanged with the stricter uncertainty threshold, suggesting these clusters may represent a reliable shape. Meanwhile, clusters one, eight, and to a lesser extent six have the highest rates of uncertainty moving from the 0.25 to 0.05 threshold, likely due to the overlap between these clusters in morphospace. The densely populated central area of morphospace may represent the generic flagellum shape with more divergent, possibly specialized shapes forming more reliable clusters in the margins of morphospace. Overall, 19% and 55% of shapes could not be clustered given a 0.25 and 0.05 threshold respectively (grey). Given this level of uncertainty, it seems that flagellum shape is a continuous character and attempts to discretize it may be subjective and misleading except in the cases of the more unique clusters that change little with a stricter uncertainty level. Coding of more specific aspects of the flagellum, as in Monjaraz-Ruedas, et al. ^12^, may ultimately give better resolution especially in areas where clusters overlap or if focusing on taxa fully within a given cluster.

Smaller genera are more likely to be exclusive to fewer clusters, with more speciose genera more frequently being found in more. For example, *Surazomus* has 27 individuals and was found in 5 clusters. Still, genera such as *Apozomus* and *Antillostenochrus* have 16 individuals classified into cluster one, while the speciose *Rowlandius* includes 52 individuals here and is found in 3 clusters. This contrasts with the correlation results of individuals per genus versus disparity, where no relationship was found. The former minimizes shape differences whereas the latter more accurately captures them. For instance, the small genus *Hubbardia* has four individuals in three clusters, giving it a relatively high ratio of individuals to clusters. However, the disparate, lanceolate shape of *Hubbardia pentapeltis* (cluster 4) is much different from the other three, resulting in a larger area of morphospace for the genus than implied by the number of clusters alone. In other words, while the number of clusters may be useful for broad categorization, it poorly captures true shape differences within genera and it is more likely that shape and how speciose a genus is are not correlated.

## Conclusion

A strength of elliptical Fourier analysis is that it only requires outlines - which all flagella have - is also a weakness, as landmarks not captured by the outline are omitted. Specific setae, setal patches, lobes, and depressions are common on male flagella, and homologizing them is an ongoing process^2^. While lobes in particular may be captured depending on the outline, other characteristics will escape morphometric quantification using this method. However, given the breadth of disparity in the male flagellum, homologization across the order may prove elusive to impossible and more specific anatomical studies of the variation in the flagellum may be more suited to genus or genus group studies.

Here, we conducted the first elliptical Fourier analysis of the arachnid order Schizomida. Examining the dorsal and lateral views in schizomids in different families, biogeographic realms, and habitats, we found support for differences between most categories examined. In more specific analyses in the Caribbean and Cuba, where there is more knowledge of total species diversity, we found similar patterns despite apparent overlaps in morphospace. Results showing differences in the amount of disparity and trends towards new areas of shape space (location) suggest that shape may be under different selective pressures based on schizomid biology, and future work may want to examine the functional repercussions of different shapes. This will likely require more in-depth work on the minutiae of shape differences and their functional importance. A well-sampled phylogeny of described species will also allow insight into rates of change, comparative evolution of shapes, and island biogeography. While clustering algorithms can cluster flagellum shapes, levels of uncertainty for some clusters suggest that the continuous nature of flagellar shape defy tidy clustering of all shapes. We hope this work may be a foundation for further study of schizomids and the utility of the male flagellum in particular to distinguish taxa in this little known lineage. Denser sampling in specific regions of interest may also aid in species delimitation, highlighting small differences in flagellum shape that may be crucial to the courtship and mating process. Last but not least, this work underscores the importance of the primary taxonomic literature to understand evolutionary patterns.

## Acknowledgments

We thank Dean Adams, Vincent Bonhomme, Chris Hamilton, Carmelo Fruciano, and Tom Stubbs for their advice and guidance improving this manuscript.

## Author contributions

The study was conceived by RJK and GSM. Data collection was done by RJK and GSM. Scripting was done by RJK and EG. Data analyses were conducted by RJK, GSM, and EG. All authors contributed to data interpretation and writing the manuscript.

## Competing interests

The authors declare no competing interests.

## Notes

https://github.com/bobkallal/schizomida

### Competing Interest Statement

The authors have declared no competing interest.

