## Supplemental File 2 for "Tales of schizomid tails: patterns in schizomid flagellum shape from elliptical Fourier analysis"

Taxonomic and image references

Abrams KM, Huey JA, Hillyer MJ, Didham RK, Harvey MS. 2020. A systematic revision of *Draculoides* (Schizomida: Hubbardiidae) of the Pilbara, Western Australia Part I: the Western Pilbara. Zootaxa 4864 DOI: 10.11646/zootaxa.4864.1.1

Armas LF. 2001. A new *Rowlandius* Reddell & Cokendolpher 1995 (Schizomida: Hubbardiidae) from Navassa Island, Greater Antilles. Rev Iber Aracnol 4:93-95.

Armas LF. 2002a. Dos generos nuevos de Hubbardiidae (Arachnida: Schizomida) de Cuba. Rev. Iber,. Aracnol. 5:3-9.

Armas LF. 2002b. Nuevas especies de *Rowlandius* Reddell & Cokendolpher, 1995 (Schizomida: Hubbardiidae) de Cuba. Rev Iber Aracnol 6:149-167.

Armas LF. 2004. Nueva especie de *Rowlandius* Reddell & Cokendolpher, 1995 (Schizomida: Hubbardiidae) de Cuba oriental. Rev Iber Aracnol 10:211-214.

Armas LF. 2007. Especie nueva de *Rowlandius* (Schizomida: Hubbardiidae) de Cuba Central. Solenodon 6: 45-51

Armas LF. 2009. Dos nuevas especies de *Hansenochrus* y *Rowlandius* (Schizomida: Hubbardiidae) de Costa Rica. Boletin Sociedad Entomologica Aragonesa 45:253-257.

Armas, LF. 2011. Genero nuevo de Hubbardiidae (Arachnida: Schizomida) para Jamaica. Solenodon 9:12-19.

Armas, LF. 2014. Two new genera of African whip scorpions (Schizomida: Hubbardiidae). Arthropoda Selecta 23(2): 97-105.

Armas LF, Abud AJ. 1990. El orden Schizomida (Arachnida) en Republica Dominicana. Poeyana 393:1-23.

Armas LF, Abud AJ. 2002. Tres especies nuevas de *Rowlandius* (Schizomida: Hubbardiidae) de Republica Dominicana, Antillas Mayores. Rev Iber Aracnol 5:11-17.

Armas LF, Cokendolpher JC. 2001. Comments on some schizomids from the Dominican Republic , with description of a new species of *Rowlandius* (Schizomida: Hubbardiidae). Rev Iber Aracnol 3:3-6.

Armas LF & Colmenares García PA. 2006. Nuevo genero de Hubbardiidae (Arachnida: Schizomida) del Zulia, Venezuela. Boletin Sociedad Entomologica Aragonesa 39:27-30.

Armas, L.F. de, and J.A. Cruz-López. 2009. Especie nueva de *Stenochrus* (Schizomida: Hubbardiidae) de Oaxaca, México. Solenodon 8: 20–24.

Armas, L.F. de & Delgado-Santa, L. (2012a) Nueva especie de *Piaroa* de la Cordillera Occidental de los Andes colombianos y segundo registro de *Stenochrus portoricensis* Chamberlin, 1922 para Colombia (Schizomida: Hubbardiidae). Boletín de la Sociedad Entomológica Aragonesa, 50, 183–186

Armas LF & Delgado-Santa L. 2012b. Nuevo genero de Hubbardiidae (Arachnida: Schizomida) de la Cordillera Occidental de los Andes, Colombia. Rev Iber Aracnol 21:139-143.

Armas LF, Villarreal OM, Colmenares-Garcia PA. 2009. Nuevo *Rowlandius* Reddell & Cokendolpher, 1995 (Schizomida: Hubbardiidae) de la Sierra San Luis, Venezuela noroccidental. Papeis Avulsos de Zoologia 49(28):361-368.

Armas, L.F. de & Víquez-Núñez, C., 2009. Primer registro del género *Piaroa* Villarreal, Giupponi et Tourinho, 2008 (Schizomida: Hubbardiidae) en Centroamérica, con la descripción de una especie nueva de Costa Rica. Boletín de la Sociedad Entomológica Aragonesa, (44), pp.131-133.

Armas LF & Rehfeldt S. 2015. *Stenochrus portoricensis*, *Zomus bagnallii* and a new genus of schizomids (Schizomida: Hubbardiidae) from a greenhouse in Frankfurt am Main, Germany. Arachnologische Mitteilungen 49: 55-61

Armas LF, Teruel R. 2002. Un genero nuevo Hubbardiidae (Arachnida: Schizomida) de las Antillas Mayores. Rev. Iber. Aracnol. 6 45-52.

Armas LF, Villarreal OM, Viquez C. 2010. Nuevas especies de *Surazomus* Reddell & Cokendolpher 1955 (Schizomida: Hubbardiidae) de Costa Rica. Papeis Avulsos de Zoologia 50.

Armas LF, Viquez C. 2010. Nuevos Hubbardiidae (Arachnida: Schizomida) de American Central. Boletín de la Sociedad Entomologica Aragonesa 46:9-21.

Armas L.F. de, Víquez C. 2011. Dos nuevas especies de *Surazomus* Reddell & Cokendolpher, 1995 (Schizomida: Hubbardiidae) de Costa Rica. Boletín de la Sociedad Entomológica Aragonesa

48: 77 –86.

Bonaldo A, Pinto-da-Rocha R. 2007. A new species of *Surazomus* (Arachnida, Schizomida) from Brazilian Oriental Amazonia. Rev. Bras. Zool. 24.

Brignoli PM. 1974. Un nuovo Schizomda delle Batu caves in Malesia. Rev. Suisse Zool. 81:731-735.

Camilo, G.R. and Cokendolpher, J.C., 1988. Schizomidae de Puerto Rico (Arachnida: chizomida). *Caribb J Sci*, *24*(1–2), pp.52-59.

Chamberlin RV. 1939. A new arachnid of the order Pedipalpia. Proc Biol Soc Washington 52:123-124.

Cokendolpher, J.C. & J.R. Reddell. 1992. Revision of the Protoschizomidae (Arachnida: Schizomida) with notes on the phylogeny of the order. Texas Memorial Museum of Speleology Monographs 3:31-74.

Cokendolpher, J. C., & Reddell, J. R. (2000). New and rare Schizomida (Arachnida: Hubbardiidae) from South America. Amazoniana: Limnologia et Oecologia Regionalis Systematis Fluminis Amazonas, 16(1-2), 187-212.

Cokendolpher J. C., D. W. Sissom and D. B. Bastawade, 1988. A new Schizomus of Indian state of Maharashtra, with additional comments on eyed schizomids (Arachnida: Schizomidae). Insecta Mundi, 2 (2): 90-96.

Cokendolpher JC, Sissom WD, Reddell JR. 2010. A new species of *Apozomus* (Arachnida: Schizomida: Hubbardiidae) from Peninsular Malaysia. Occasional Papers, Museum of Texas Tech University 298.

Cook 1899. *Hubbardia*, a new genus of Pedipalpi. Proceedings of the Entomological Society of Washington, vol. 4, p. 249-261.

Delgado-Santa L, Armas LF. 2013. Tres nuevos Hubbardiinae (Schizomida: Hubbardiidae) de Colombia. Rev. Iber. Aracnol. 22:37-45.

Dumitresco M. 1973. Deux espèces nouvelles du genre *Schizomus* (Schizomida), trouvées aCuba. Resultats des expéditions biospéléologiques cubano-rumaines aCuba, 1, 279-292.

Dumitresco M. 1977. Autres nouvelles espèces du genre *Schizomus* des grottes de Cuba // Résultats des expéditions biospéologiques cubano-roumaines à Cuba. Bucuresti: Editura Academiei. Vol.2.

P.147–158.

Gertsch, W. J. 1940. Two new American whip-scorpions of the family Schizomidae. American Museum Novitates, 1077, 4pp.

Giupponi APL, Miranda GS, Villarreal OM (2016) *Rowlandius dumitrescoae* species group: new diagnosis, key and description of new cave-dwelling species from Brazil (Schizomida, Hubbardiidae). ZooKeys 632: 13–34. doi: 10.3897/zookeys.632.9337

González-Sponga MA. 1997. Arácnidos de Venezuela: Un nuevo género y dos nuevas especies de Schizomidae y redescripción de *Schizomus* simoni Hansen y Sorensen, 1995 del sistema montañoso de la costa (Schizomida). Acta Biológica Venezuelica, vol. 17, no 2, p. 1-10.

Hansen & Sørensen, 1905 : The Tartarides, a tribe of the order Pedipalpi. Arkiv för Zoologi, vol. 2, no 8, p. 1-78

Hansen, H.J. (1921). The Pedipalpi, Ricinulei, and Opiliones (exc. Op. Laniatores) collected by Mr.

Leonardo Fea in tropical West Africa and adjacent islands. In, Studies on Arthropoda, vol. 1: 1–55. Gyldendalske Boghandel: Kjøbenhavn.

Harvey, M. S. (1988). A new troglobitic schizomid from Cape Range, Western Australia (Chelicerata: Schizomida). Records of the Western Australian Museum 14, 15–20.

Harvey MS. 1992. The Schizomida (Chelicerata) of Australia. Invertebrate Taxonomy 6(1):77-129.

Harvey MS. 2000. A review of the Australian schizomid genus *Notozomus* (Hubbardiidae). Memoirs of the Queensland Museum 46: 161-174.

Harvey MS. 2000. *Brignolizomus* and *Attenuizomus*, new schizomid genera from Australia (Schizomida Hubbardiidae). Mem Soc Entomol Ital 78(2):329-338.

Harvey MS. 2001b. The Schizomida (Arachnida) of the Seychelle Islands. Invertebrate Taxonomy 15(5):681-693.

Harvey MS. 2001c. New cave-dwelling schizomids (Schizomida: Hubbardiidae) from Australia. Records of the Western Australian Museum 64:171-185.

Harvey MS. 2006. The schizomid fauna (Arachnida: Schizomida: Hubbardiiidae) of the Arabian Peninsula and Somalia. Fauna of Arabia. 21: 167-177.

Harvey MS, Abrams KM. 2015. A new troglobitic schizomid (Hubbardiidae: *Paradraculoides*) from the Pilbara region, Western Australia. Records of the Western Australian Museum 30: 132-136.

Harvey MS, Berry O, Edward KL, Humphreys G. 2008. Molecular and morphological systematics of hypogean schizomids (Schizomida: Hubbardiidae) in semiarid Australia. Invert Syst 22(2):167-194.

Harvey MS & Humpreys WF. 1995. Notes on the genus *Draculoides* Harvey (Schizomida: Hubbardiidae), with the description of a new troglobitic species. Records of the Western Australian Museum 52:183-189.

Kraus O. 1957. Schizomidae aus Kolumbien (Arach., Pedipalpi-Schizopeltidia). Senckenb Biol. 38: 245–250.

Kraus O, Beck L. 1967. Taxonomie und Biologie von *Trithyreus brasiliensis* n. sp. (Arach.: Pedipalpi: Schizopeltidia). Senckenberg Biol. 48:401-405.

Kishida, K. 1930. On the occurrence of the genus *Trithyreus* in Bonin Islands. Lansania, Tokyo, 2: 17-19

Lawrence RF. 1958. Whipscorpions (Uropygi) from Angola, the Belgian Congo and Mossambique. Companhia de Diamantes de Angola, vol. 40. Publicacões Culturais, pp. 69–79.

Lawrence RF. 1969. The Uropygi (Arachnida: Schizomidae) of the Ethiopidan region. Journal of Natural History 3:217-260.

McDonald WA, Hoge CL. 1957. A new *Trithyreus* from Southern California (Pedipalpi, Schizomidae). American Museum Novitates 1834:1-8.

Mello-Leitão, C. (1931). Pedipalpos do Brasil e algumas notas sobre a ordem. Archivos do Museu

Nacional 33: 7–72.

Monjaraz-Ruedas, R. 2012. A new species of the schizomid genus *Stenochrus* (Schizomida: Hubbardiidae) from Mexico. Zootaxa 68 (3334): 63–68.

Monjaraz-Ruedas R. 2013. A new species of *Protoschizomus* (Schizomida: Protoschizomidae) from a cave in Guerrero, Mexico. JoA 41(3):420-424.

Monjaraz-Ruedas R, Francke OF 2015. Taxonomic revision of the genus *Mayazomus* Reddell & Cokendolpher, 1995 (Schizomida: Hubbardiidae), with description of five new species from Chiapas, Mexico. Zootaxa 3915: 451–490.

Monjaraz-Ruedas R & Francke OF. 2017. A new genus of schizomids (Arachnida: Schizomida: Hubbardiidae) from Mexico, with notes on its systematics. Systematics and Biodiversity 2000: 1–15.

Monjaraz-Rueas R, Francke OF. 2018. Five new species of *Stenochrus* (Schizomida: Hubbardiidae) from Oaxaca, Mexico. Zootaxa 4374:2.

Monjaraz-Ruedas R, Francke OF, Cokendolpher J. 2016. Three new species of *Agastoschizomus* (Arachnida: Schizomida: Protoschizomidae) from North America. Revista Mexicana de Biodiversidad. 87:16. 10.1016/j.rmb.2016.02.006.

Monjaraz-Ruedas R, Prendini L, Francke OF. 2019. Systematics of the Short-Tailed Whipscorpion Genus *Stenochrus* Chamberlin, 1922 (Schizomida: Hubbardiidae), with Descriptions of Six New Genera and Five New Species. 435:1-191. doi.org/10.1206/0003-0090.435.1.1

Monjaraz-Ruedas R, Prendini L, Francke OF. 2020. First species of *Surazomus* (Schizomida: Hubbardiidae) from North America illuminate biogeography of short-tailed whipscorpions in the New World. Arthropod Systematics & Phylogeny 78(2):245-263.

Moreno HM & Francke OF. 2009. A new species of Agastoschizomus (Schizomida: Protoschizomidae) from Guerrero, Mexico. Texas Memorial Museum Speleological Monographs, 7. Studies on the cave and endogean fauna of North America, V. Pp. 33-36.

Moreno-González JA, Villarreal MO. 2012. A new genus of Hubbardiidae (Arachnida: Schizomida) from the Colombian Andes, with some taxonomic comments. Zootaxa. 3560:61–78

Moreno-González JA & Villarreal-M O. (2017) Two new species of *Calima* Moreno-González and Villarreal, 2012 (Arachnida: Schizomida: Hubbardiidae) from the Colombian Andes, with a discussion on the male flagellar microsetae of Hubbardiinae, Journal of Natural History, 51:45-46, 2681-2700, DOI: 10.1080/00222933.2017.1397226

Moreno-Gonzalez JA, Delgado-Santa L, Armas LF. 2009. Two new species of *Piaroa* (Arachnida: Schizomida, Hubbardiidae) from Colombia, with comments on the genus taxonomy and the flagellar setae pattern of Hubbardiindae. Zootaxa 3852 (2): 227–251

Pinto-da-Rocha R. 1996. *Surazomus chavin* new species, first Schizomida (Hubbardiidae, Hubbardiinae) described from Peru. JoA 24(3):265-267.

Pinto-da-Rocha R, Andrade R & Moreno-González JA. 2016. Two new cave-dwelling genera of short-tailed whip-scorpions from Brazil (Arachnida: Schizomida: Hubbardiidae). Zoologia 33, No. 2, pp. 1-9.

Reddell JR, Cokendolpher JC. 1984. A new species of troglobitic *Schizomus* (Arachnida: Schizomida) from Ecuador. Bull Br Arachnol Soc. 6: 172‒177.

Reddell, J.R. & Cokendolpher, J.C. (1986) New species and records of *Schizomus* (Arachnida: Schizomida) from Mexico. Texas Memorial Museum, Speleological Monographs, 1, 31–38.

Reddell, J.R. and Cokendolpher, J.C. (1991). Redescription of *Schizomus* *crassicaudatus* (Pickard-

Cambridge) and diagnoses of *Hubbardia* Cook, *Stenochrus* Chamberlin, and *Sotanostenochrus*

new genus, with description of a new species of *Hubbardia* from California (Arachnida: Schizomida:

Hubbardiidae). Pearce Sellards Series 47:1–24.

Reddell, James R., and James C. Cokendolpher. 1995. "Speleological Monographs, no. 4. Catalogue, bibliography, and generic revision of the order Schizomida (Arachnida). Catálogo, bibliografía y revisión genérica del orden Schizomida (Arachnida)."

Remy, P. 1961. Sur l'ecologie des Schizomides (Arachn. Uropyges) de mes recoltes, avec description de trois Schizomus nouveaux, captures par J. van der Drift au Surinam. Bull. Mus. Natl. Hist. Nat., Ser. 2, 33(4):406-414, 500-511.

Rowland, J. M. (1971). *Agastoschizomus lucifer*, a new genus and species of cavernicole schizomid (Arachnida, Schizomida) from Mexico. Bulletin of the Association for Mexican Cave Studies 4, 13-17.

Rowland, J.M. 1973a. A new genus and several new species of Mexican schizomids (Schizomida: Arachnida). Occasional Papers of the Museum, Texas Tech University 11: 1–23.

Rowland, J.M. 1973c. Three new Schizomida of the genus *Schizomus* from Mexican caves (Arachnida). Bulletin of the Association for Mexican Cave Studies 5: 135–140.

Rowland, J.M. 1975. A partial revision of Schizomida (Arachnida), with descriptions of new species, genus, and family. Occasional Papers of the Museum, Texas Tech University 31:1-21.

Rowland, J.M. and Reddell, J.R., 1977. A review of the cavernicole Schizomida (Arachnida) of Mexico, Guatemala, and Belize. Assoc. Mexican Cave Stud. Bull, 6, pp.79-102.

Rowland JM, Reddell JR. 1979. The order Schizomida (Arachnida) in the New World I. Protoschizomidae and *Dumitrescoae* group (Schizomidae: *Schizomus*). JoA 6:161-196.

Rowland JM, Reddell JR. 1980. The order Schizomida (Arachnida) in the New World III. Mexicanus and pecki groups (Schizomidae: *Schizomus*). JoA 8:1-34.

Rowland JM, Reddell JR. 1981. The order Schizomida (Arachnida) in the New World IV. Goodnightorum and briggsi groups and unplaced species. JoA 9: 19-46.

Ruiz GRS, Valente RM. 2017. The first schizomid from a dry forest in South America (Arachnida: Schizomida). Zootaxa DOI: 10.11646/ZOOTAXA.4311.1.5

Ruiz GRS, Valente RM. 2019. Description of a new species of *Surazomus* (Arachnida: Schizomida), with comments on homology of male flagellum and mating march anchorage in the genus. Plos One

10.1371/journal.pone.0213268

Salvatierra L. 2018. A new species of *Surazomus* Reddell and Cokendolpher 1995 (Arachnida: Schizomida) from Rondonia, Brazil. Turkish Journal of Zoology 42:107-112.

Santos AJ, Dias SC, Brescovit AD, Santos PP. 2008. The arachnid order Schizomida in the Brazilian Atlantic Forest: a new species of *Rowlandius* and new records of *Stenochrus portoricensis* (Schizomida: Hubbardiidae). Zootaxa 1850:53-60.

Santos AJ, Ferreira RL, Buzzato BA. 2013. Two new cave-dwelling species of the short-tailed whipscorpion genus *Rowlandius* (Arachnida: Schizomida: Hubbardiidae) from northeastern Brazil, with comments on male dimorphism. PlosOne 8(5):e63616. https://doi.org/10.1371/journal.pone.0063616

Santos A, Pinto-da-Rocha R. 2009. A new micro-whip scorpion species from Brazilian Amazonia (Arachnida, Schizomida, Hubbardiidae), with the description of a new synapomorphy for Uropygi. JoA 37:39-44.

Segovia-Paccini A, Ahumada-C D, Moreno-Gonzalez JA. 2018. A new remarkable short-tailed whip-scorpion species of *Piaroa* (Arachnida, Schizomida, Hubbardiidae) from the Colombian Caribbean region. Zootaxa 4500:91-103.

Sekiguchi K, Yamasaki T. 1972. A redescription of “*Trithyreus sawadai*” (Uropygi: Schizomidae) from the Bonin Islands. Acta Arachnologica 24: 73-81.

Sissom WD. 1980. The eyed schizomids, with a description of a new species from Sumatra (Schizomida: Schizomdae). JoA 8(2):187-192.

Teruel R. 2000. Una nueva especie de *Rowlandius* Reddell & Cokendolpher, 1995 (Schizomida: Hubbardia) de Cuba Oriental. Rev. Iber. Aracnol. Vol.1, XII pp: 45–47.

Teruel R. 2003. Adiciones a la fauna cubana de esquizómidos, con la descripción de un nuevo género y nueve especies nuevas de Hubbardiidae. Rev. Iber. Aracnol. 7 39-69

Teruel R. 2004. Nuevas adiciones a la fauna de esquizómidos de Cuba oriental, con la descripción de cuatro nuevas especies (Schizomida: Hubbardiidae). Rev. Iber. Aracnol. 9 31-42

Teruel, R. 2007. Los esquizómidos troglomorfos de Cuba, con las descripciones de dos géneros y una especie nuevos (Schizomida: Hubbardiidae: Hubbardiinae). Boletín de la Sociedad Entomológica Aragonesa, 40: 39-53.

Teruel R. 2012. Un nuevo *Rowlandius* Reddell & Cokendolpher 1995 del macizo de Guamuhaya, Cuba central (Schizomida: Hubbardiidae). Rev Iber Aracnol 21:61-64.

Teruel R. 2013. Un nuevo *Antillostenochrus* Armas & Teruel 2002 de Cuba centro-oriental (Schizomida: Hubbardiidae). Rev. Iber. Aracnol. 22 61-65.

Teruel R. 2015. Una especie nueva de *Antillostenochrus* Armas & Teruel 2002 (Schizomida: Hubbardiidae), del extremo oriental de Cuba. Revista Iberica de Aracnologia 27:75-80.

Teruel, R. (2017). Revisión taxonómica del género *Cubazomus* Reddell & Cokendolpher, 1995, con la descripción de una especie nueva de Cuba (Schizomida: Hubbardiidae). Revista Ibérica de Aracnología, 30, 71-81.

Teruel R. 2018. Two new genera and a new species of schizomids (Arachnida: Schizomida) from Isla de Pinos, Cuba. Ecologica Montenegrina 19:33-49

Teruel R, Armas LF. 2002. Un género nuevo de Hubbardiidae (Arachnida: Schizomida) del occidente de Cuba. Revista Iberica de Aracnologia 6:91-94.

Teruel R, Armas LF. 2004. Un nuevo *Rowlandius* Reddell & Cokendolpher 1995 de la Sierra Maestra, Cuba oriental (Schizomida: Hubbardiidae). Rev Iber Aracnol 21:5-8.

Teruel R, Armas LF, Rodriguez TM. 2012. Adiciones a los esquizómidos de Cuba central, con la descripción de cuatro nuevos *Rowlandius* Reddell & Cokendolpher 1995 (Schizomida: Hubbardiidae). Rev Iber Aracnol 21:97-112.

Teruel R, Rodriguez-Cabrera TM. 2019. Two remarkable species of Hubbardiidae Cook, 1899 (Arachnida: Schizomida) from eastern Cuba. Ecologica Montenegrina 20:40-54.

Villarreal OM & García, L.F., 2012. A new species of *Piaroa* Villarreal, Giupponi & Tourinho, 2008 (Schizomida: Hubbardiidae) from Colombia. Turkish Journal of Zoology, 36(2), pp.185-189.

Villarreal OM, Armas LF, Garcia LF. 2014. A new species of *Piaroa* (Schizomida: Hubbardiidae) from Venezuela, with taxonomic notes on the genus. Zootaxa 3765:371-381.

Villarreal OM, Teruel R. 2006. Un nuevo *Hansenochrus* Reddell & Cokendolpher, 1995 (Schizomida: Hubbardiidae) de Venezuela noroccidental. Papers Avulsos de Zoologia 46(20):233-238.

Villarreal OM, Giupponi APL, Tourinho AL.2008. New Venezuelan genus of the Hubbardiidae (Arachnida: Schizomida). Zootaxa 1860:60-68.

Villarreal OM, Miranda GS, Giupponi APL. 2016. New proposal of setal homology in Schizomida and revision of *Surazomus* from Ecuador. Plos One doi: 10.1371/journal.pone.00147012
