## Supplementary figures and images for "Tales of schizomid tails: patterns in schizomid flagellum shape from elliptical Fourier analysis"

### Han_vanderdrifti.jpg

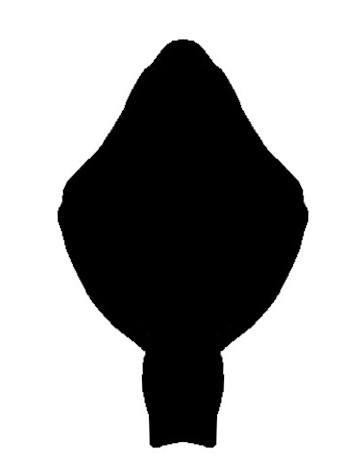

### Han_yolandae.jpg

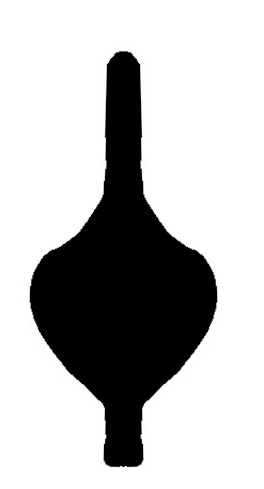

### Har_contrerasi.jpg

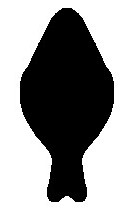

### Har_mexicanus.jpg

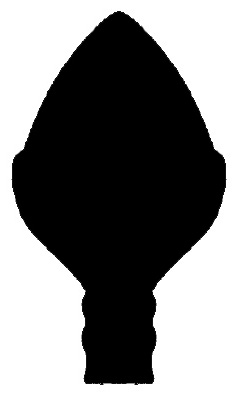

### Har_mulaiki.jpg

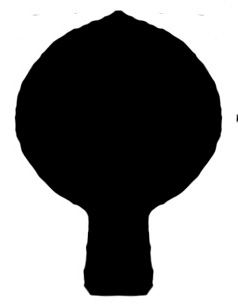

### Har_reddelli.jpg

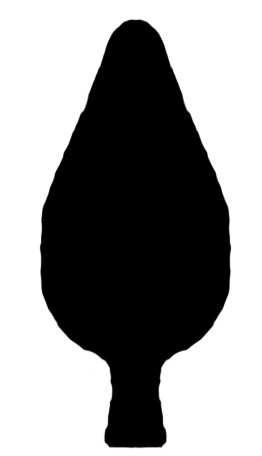

### Het_goodnightorum.jpg

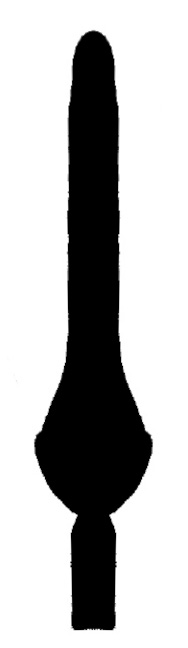

### Het_kekchi.jpg

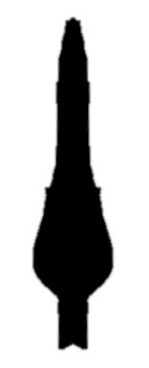

### Het_meambar.jpg

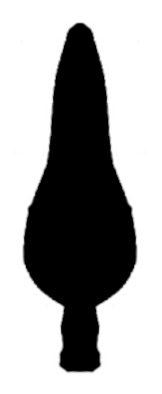

### Het_orthoplax.jpg

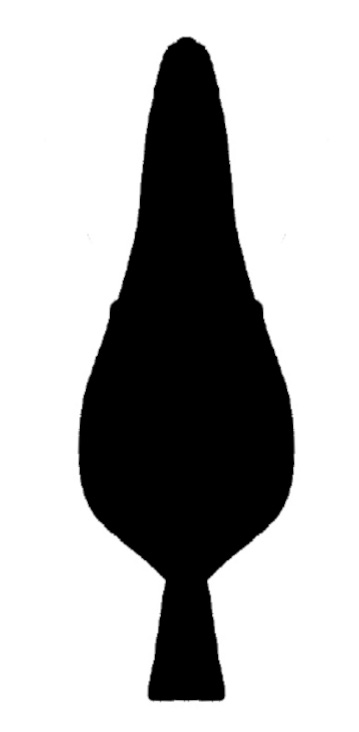

### Het_silvino.jpg

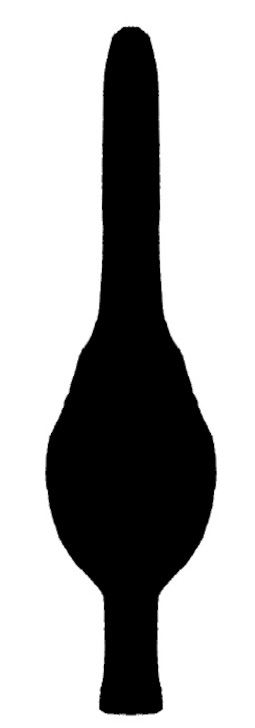

### Htc_sierramaestrae.jpg

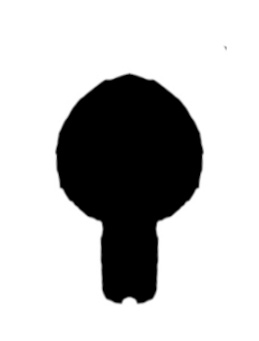

### Hub_belkini.jpg

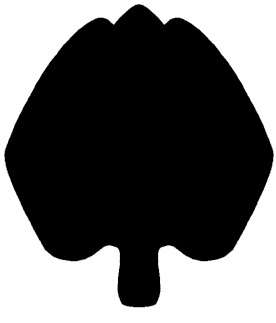

### Hub_joshuaensis.jpg

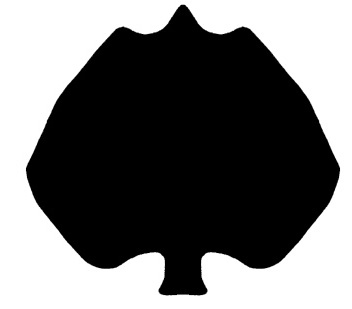

### Hub_pentapeltis.jpg

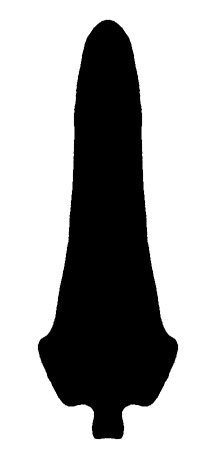

### Hub_wessoni.jpg

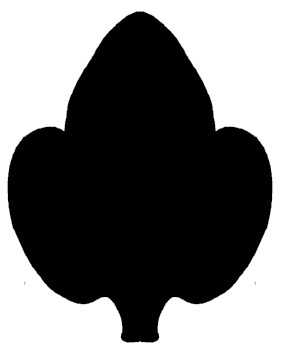

### Jul_cooloola.jpg

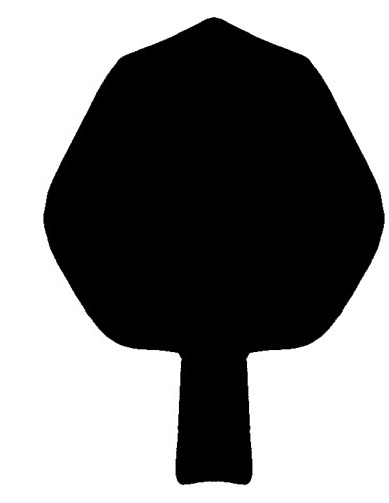

### Jul_lawrencei.jpg

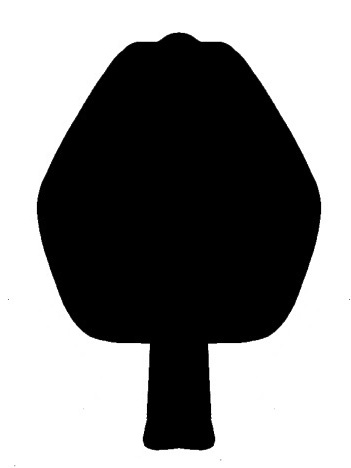

### Ken_pekkai.jpg

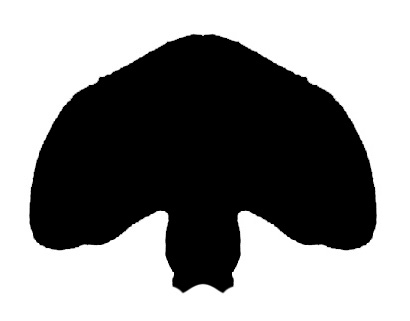

### Law_atlanticus.jpg

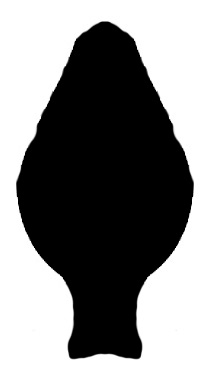

### Law_bong.jpg

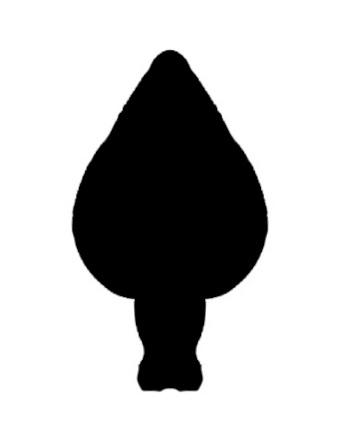

### Lui_yunquensis.jpg

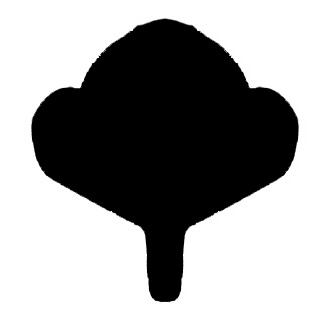

### May_aluxe.jpg

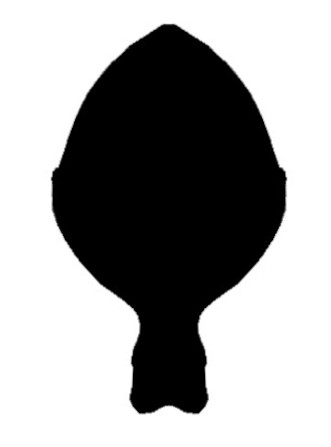

### May_estorAV10.jpg

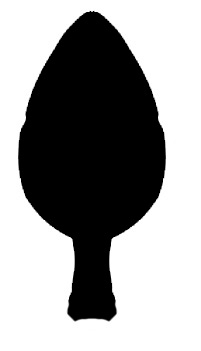

### May_estorMR16.jpg

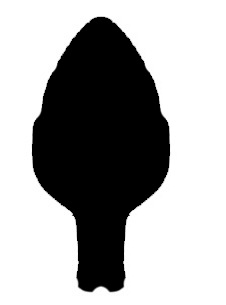

### May_hoffmannae.jpg

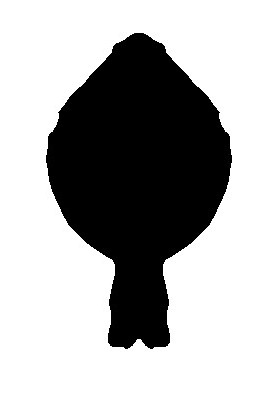

### May_infernalis.jpg

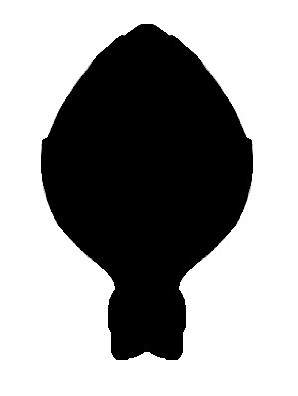

### May_kaamuul.jpg

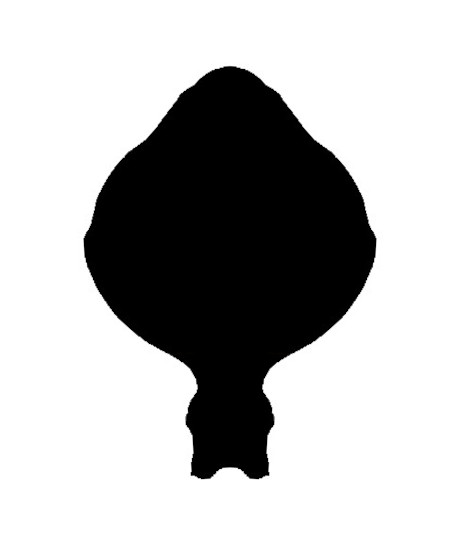

### May_loobil.jpg

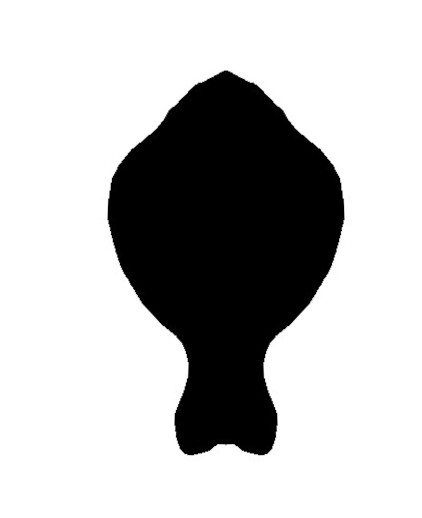

### May_tzotzil.jpg

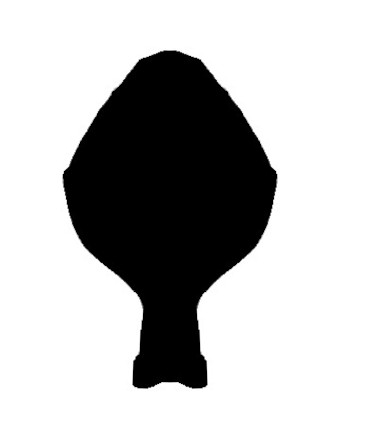

### May_yaax.jpg

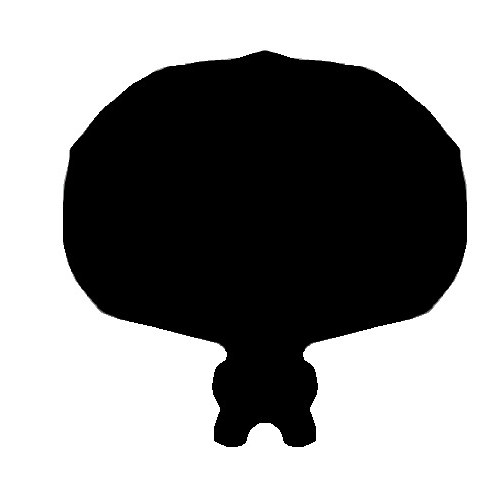

### Meg_mossambicus.jpg

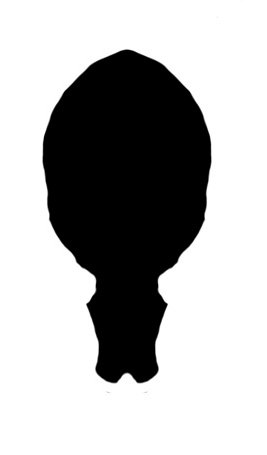

### Nad_carajas.jpg

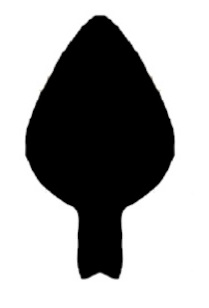

### Nah_bokmai.JPG

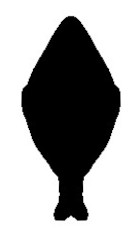

### Nah_caballero.jpg

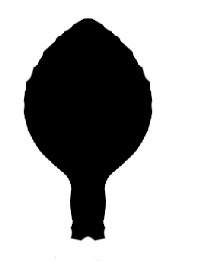

### Nah_lanceolatus.jpg

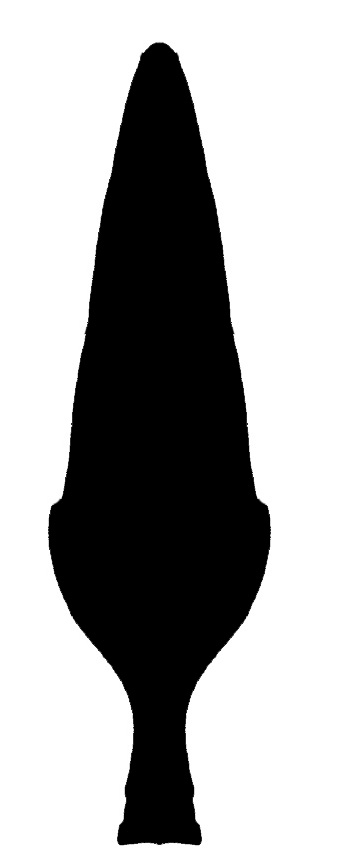

### Nah_pallidus.jpg

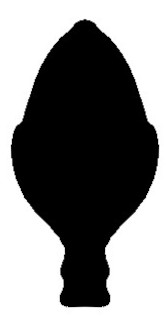

### Neo_tikaderi.jpg

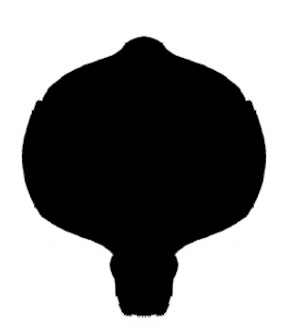

### Not_aterpes.jpg

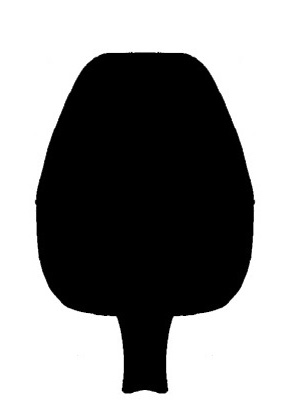

### Not_bronwenae.jpg

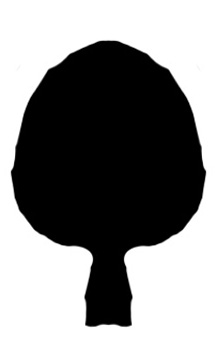

### Not_curiosus.jpg

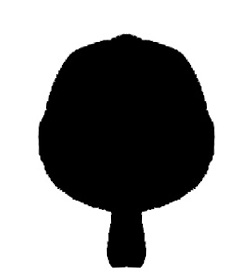

### Not_daviesae.jpg

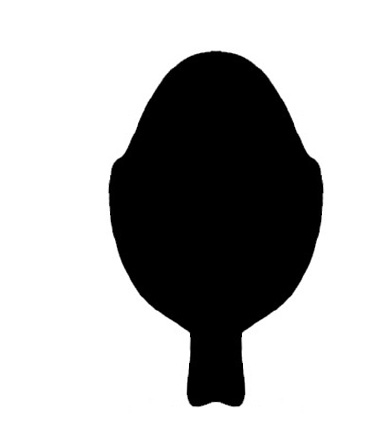

### Not_ingham.jpg

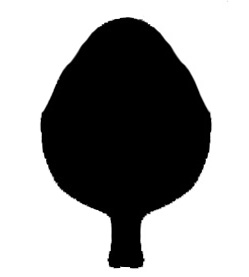

### Not_jacquelinae.jpg

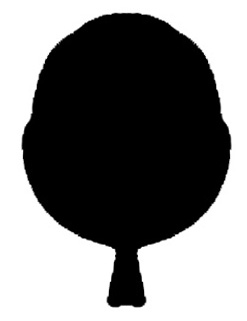

### Not_ker.jpg

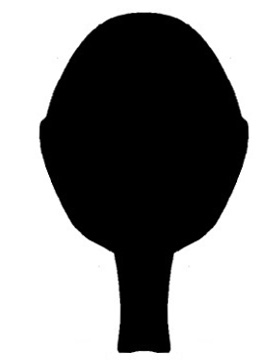

### Not_majesticus.jpg

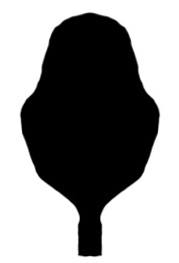

### Not_maurophila.jpg

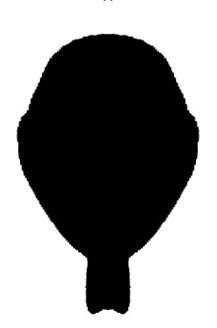

### Not_rentzi.jpg

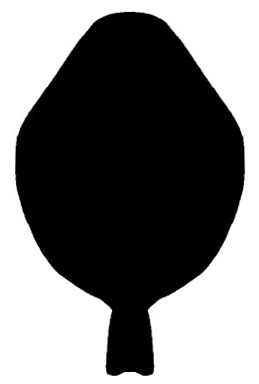

### Not_spec.jpg

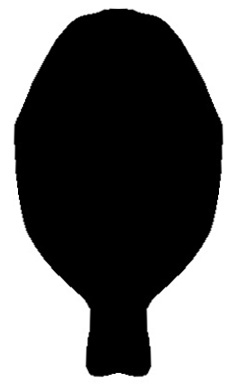

### Not_wudjl.jpg

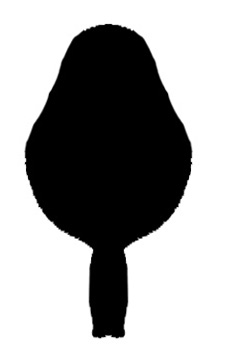

### Ocu_biocellatus.jpg

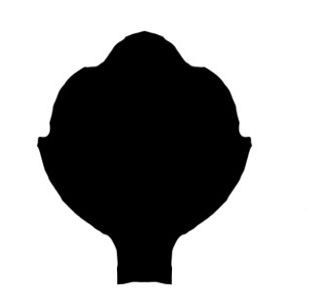

### Olm_brujo.jpg

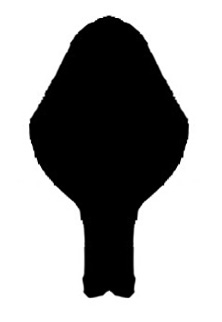

### Olm_cruzlopezi.jpg

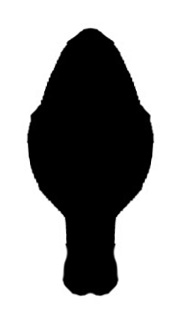

### Olm_santibanezi.jpg

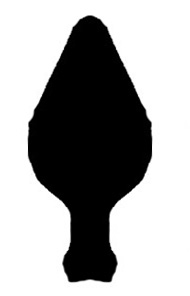

### Ori_luzonicus.jpg

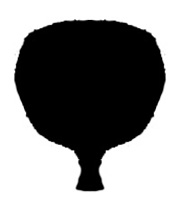

### Ori_sawadai.jpg

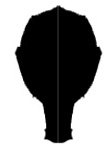

### Ovo_lunatus.jpg

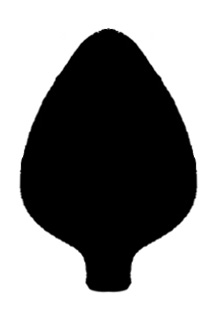

### Pac_lacandonus.jpg

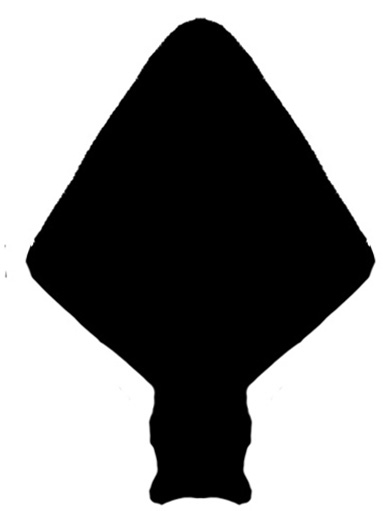

### Pac_moisii.jpg

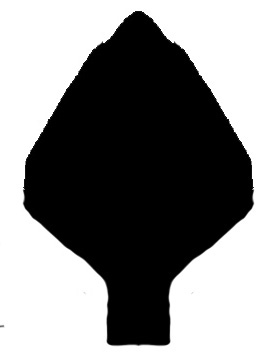

### Pac_stewarti.jpg

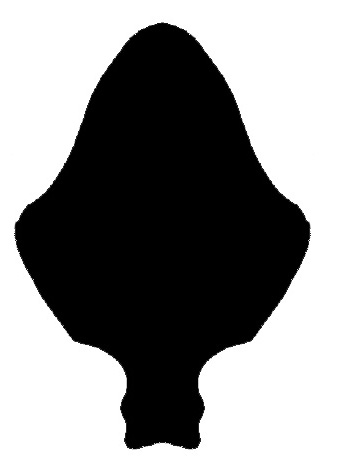

### Pac_tepezcuintle.jpg

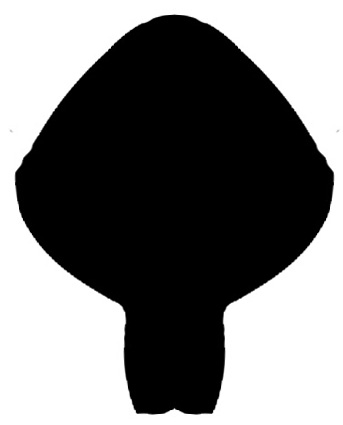

### Pac_trilobatus.jpg

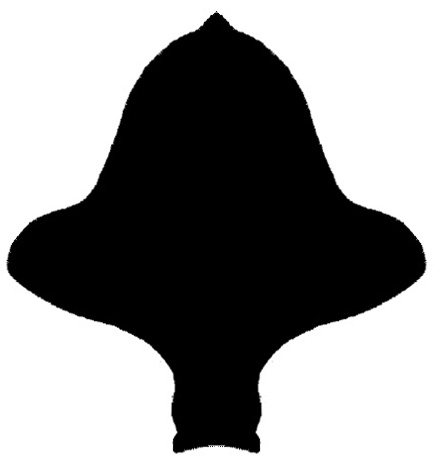

### Pia_bacata.jpg

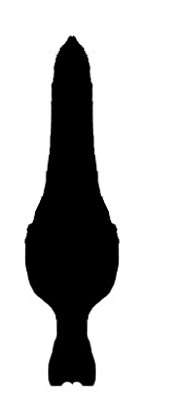

### Pia_bijagua.jpg

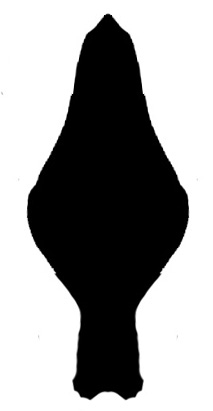

### Pia_escalerete.jpg

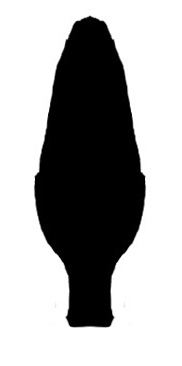

### Pia_guipongai.jpg

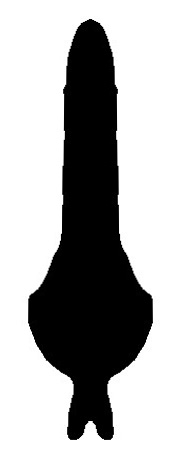

### Pia_pioi.jpg

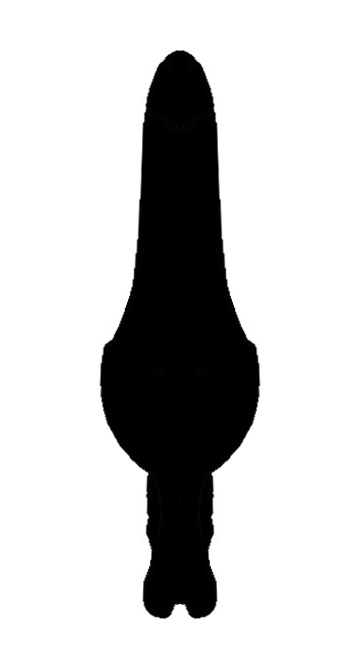

### Pia_turbacoensis.jpg

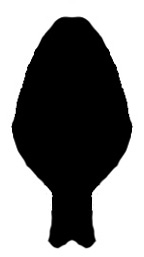

### Pia_villarreali.jpg

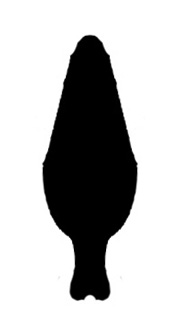

### Pia_virichaj.jpg

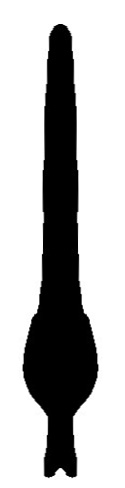

### Pia_youngi.jpg

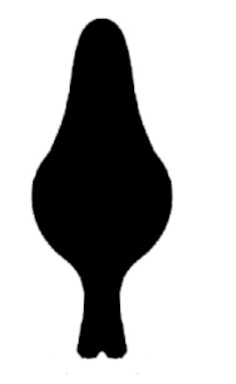

### Pin_marmoreus.jpg

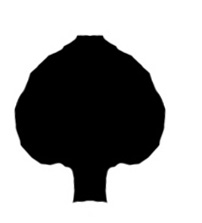

### Pro_franckei.jpg

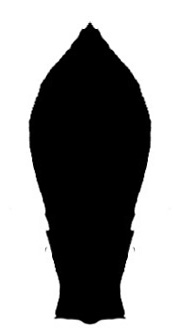

### Pro_occidentalis.jpg

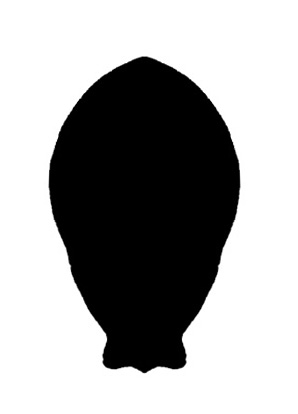

### Pro_pachypalpus.jpg

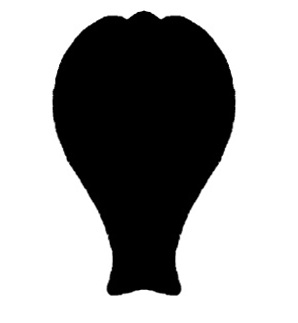

### Pro_rowlandi.jpg

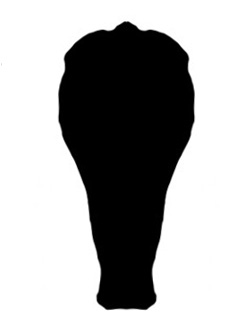

### Pro_sprousei1.jpg

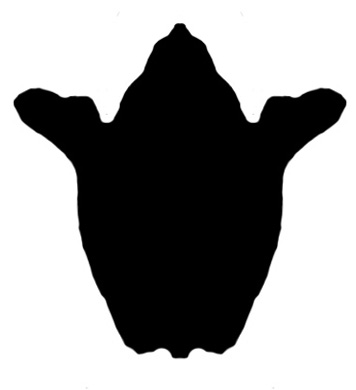

### Pro_sprousei2.jpg

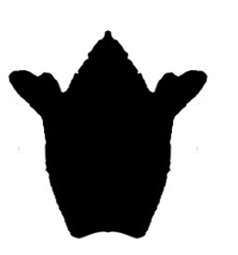

### Red_cubensisArmas.jpg

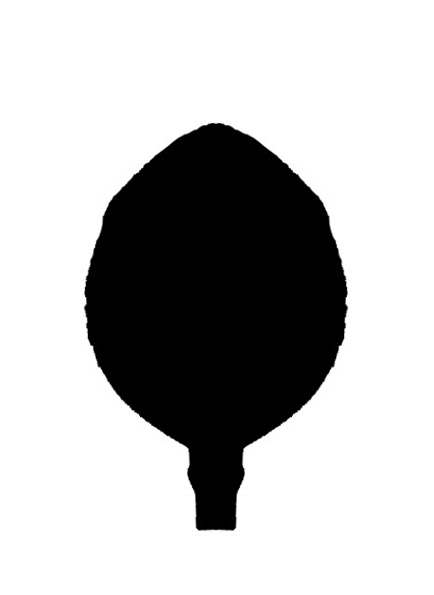

### Red_cubensisTer.jpg

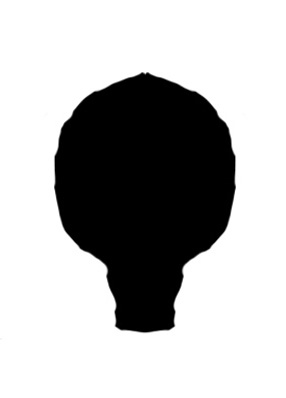

### Row_abeli.jpg

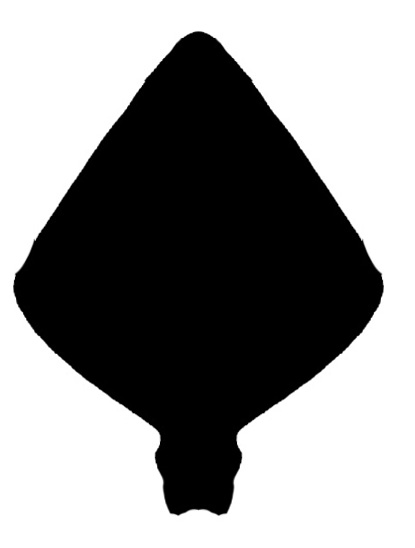

### Row_anasilviae.jpg

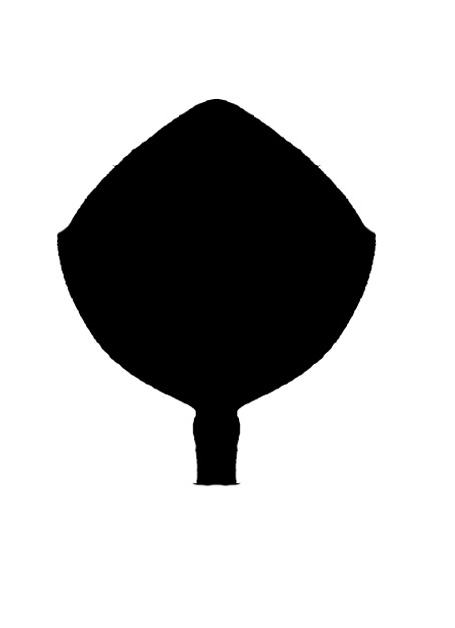

### Row_arduus.jpg

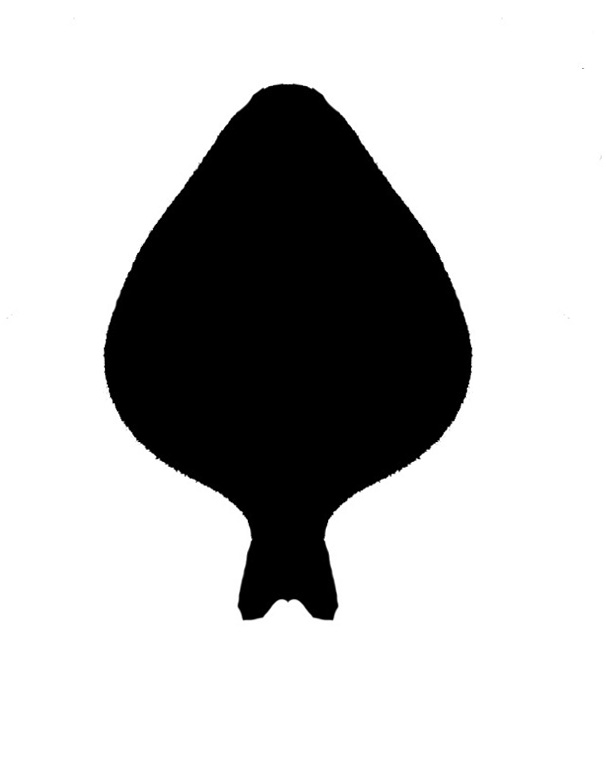

### Row_arenicola.jpg

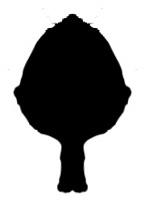

### Row_biconourus.jpg

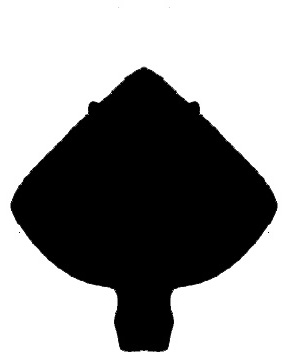

### Row_candidae.jpg

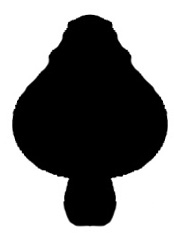

### Row_casabito.jpg

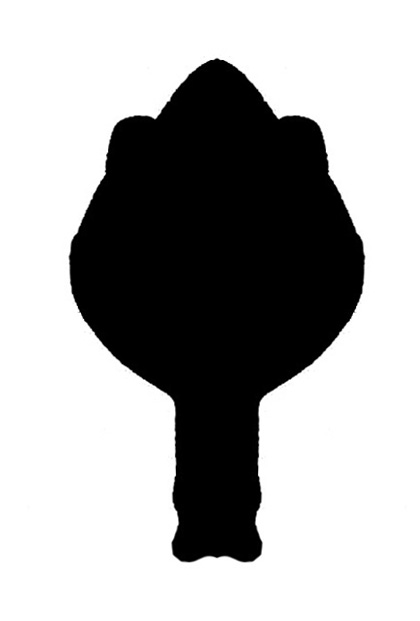

### Row_cousinensis.jpg

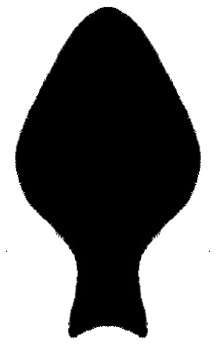

### Row_cupeyalensis.jpg

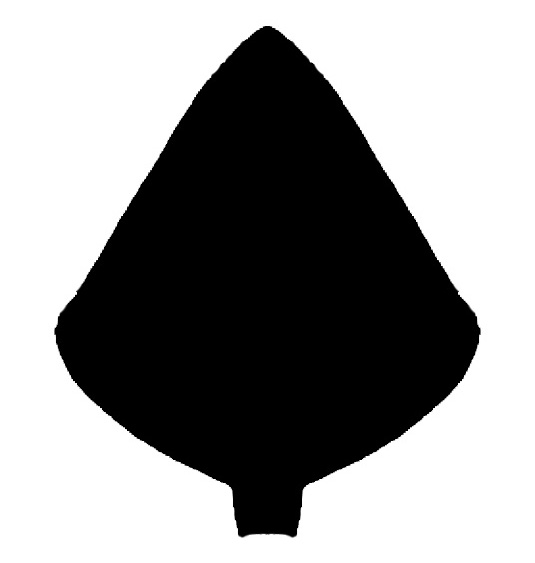

### Row_desecheo.jpg

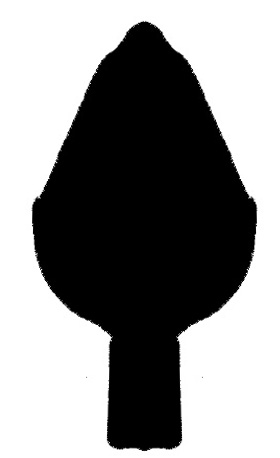

### Row_ducoudrayi.jpg

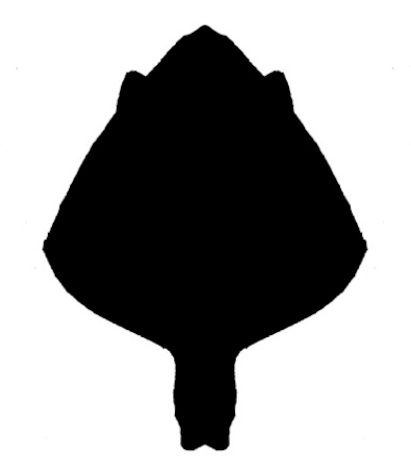

### Row_dumitrescoae.jpg

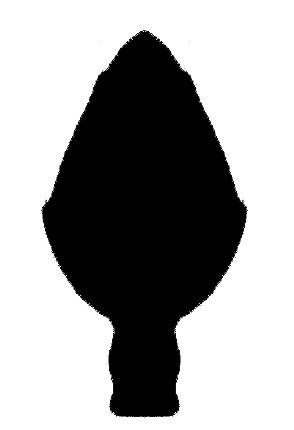

### Row_engombe.jpg

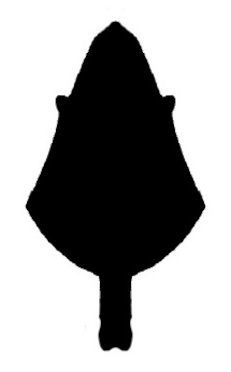

### Row_falcifemur.jpg

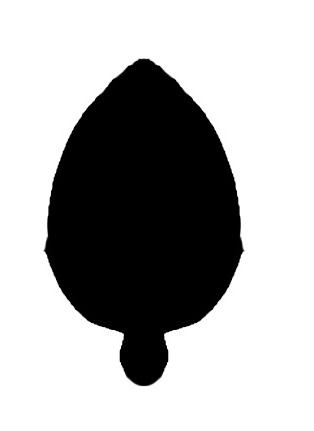

### Row_florenciae.jpg

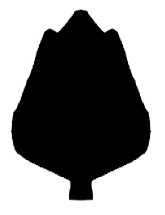

### Row_gladiger.jpg

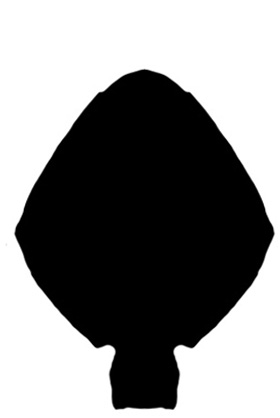

### Row_gracilis.jpg

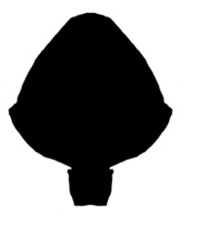

### Row_guama.jpg

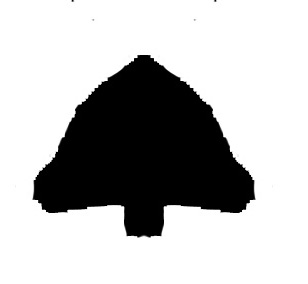

### Row_guamuhaya.jpg

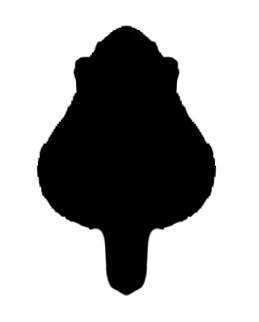

### Row_guantanamero.jpg

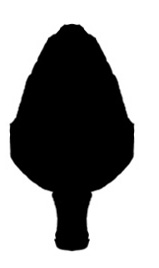
