## Supplementary figures and images for "Tales of schizomid tails: patterns in schizomid flagellum shape from elliptical Fourier analysis"

### Adi_duckei.jpg

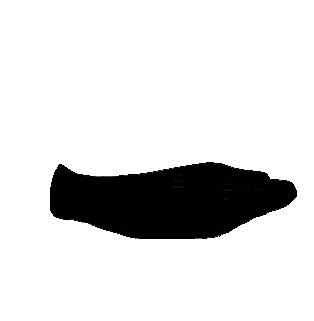

### Aga_huitzmolotitlensis.jpg

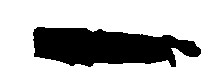

### Aga_juxtlahuacensis.jpg

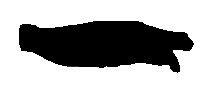

### Aga_lucifer.JPG

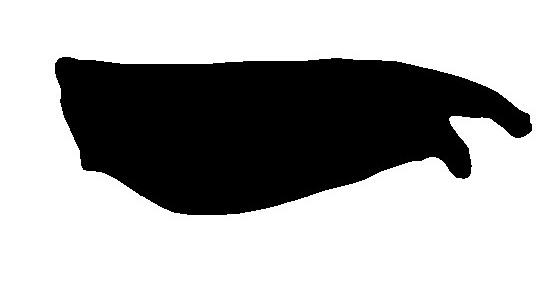

### Aga_patei.jpg

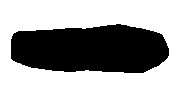

### Aga_tamaulipensis.jpg

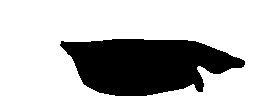

### Amb_aquismon.jpg

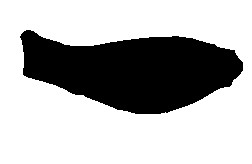

### Amb_davisi.JPG

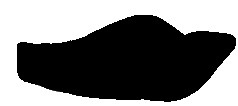

### Amb_montielae.jpg

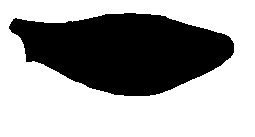

### Ane_harteni.jpg

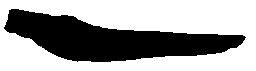

### Ant_anseli.jpg

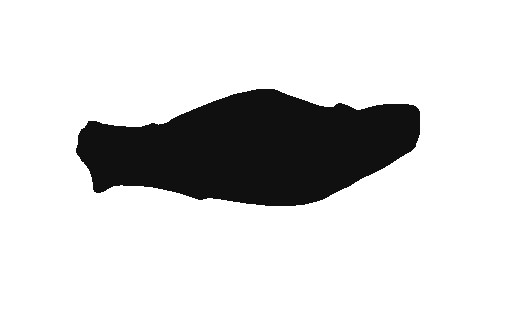

### Ant_brevipatellatus.jpg

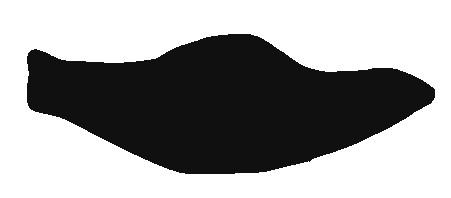

### Ant_cokendolpheri.jpg

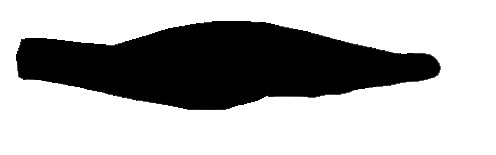

### Ant_eremita.jpg

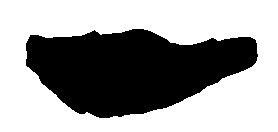

### Ant_gibarensis.jpg

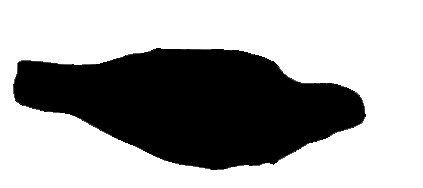

### Ant_holguin.jpg

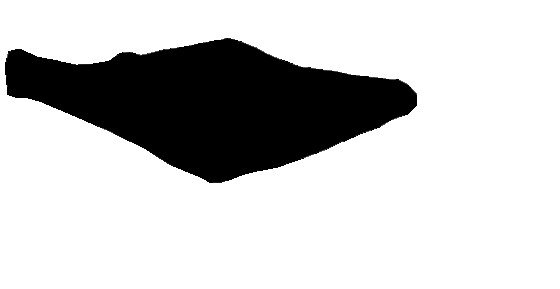

### Ant_longior.jpg

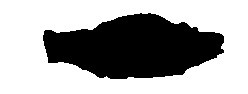

### Ant_planicauda.jpg

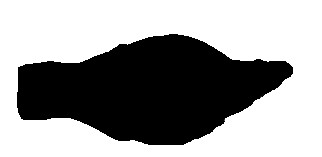

### Apo_alligator.jpeg

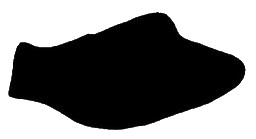

### Apo_brignolii.jpeg

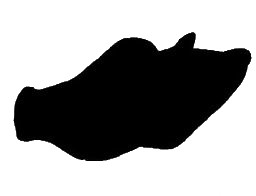

### Apo_eberhardi.JPG

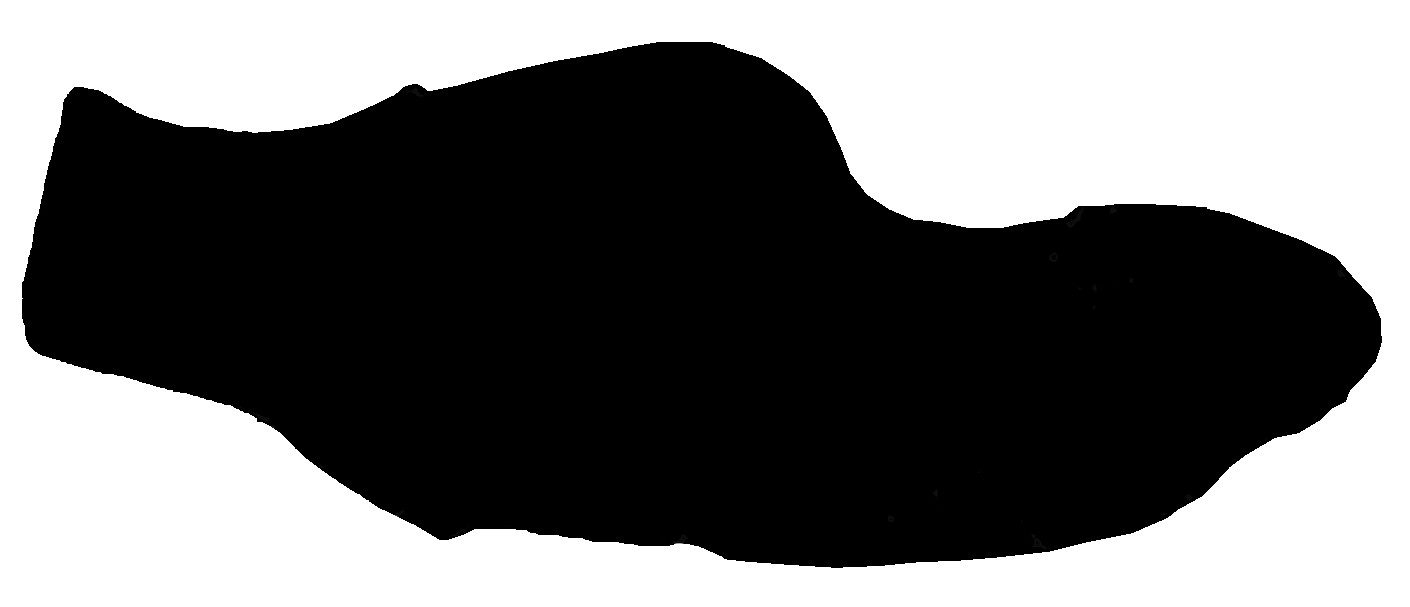

### Apo_gerlachi.jpg

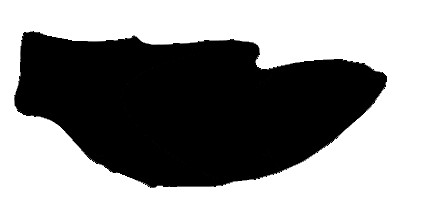

### Apo_howarthi.jpg

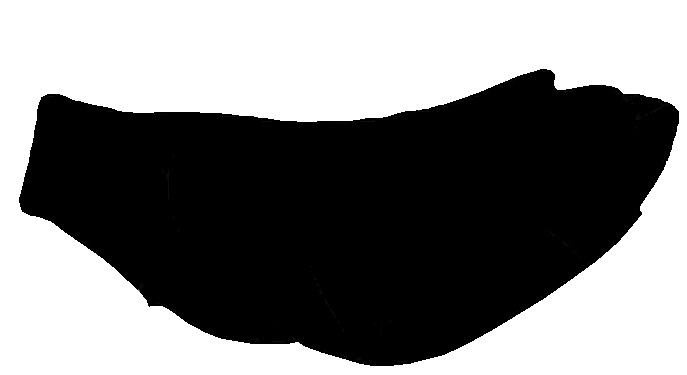

### Apo_termitarium.jpg

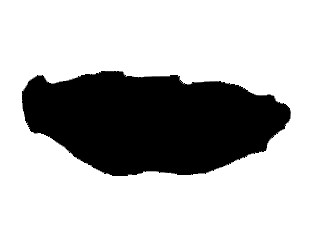

### Apo_watsoni.jpeg

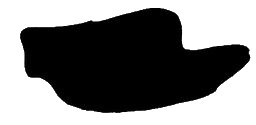

### Apo_yirrkala.jpeg

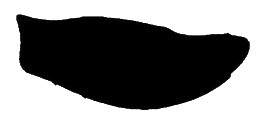

### Art_liberiensis.JPG

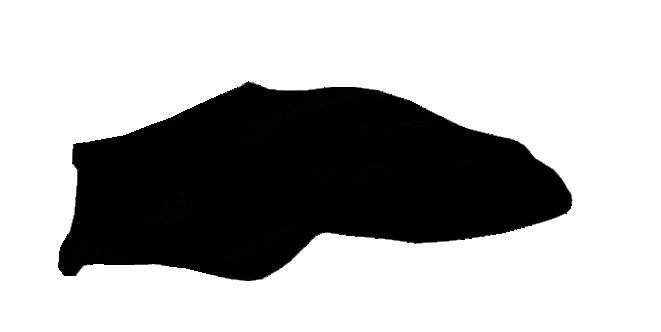

### Att_baroalba.jpg

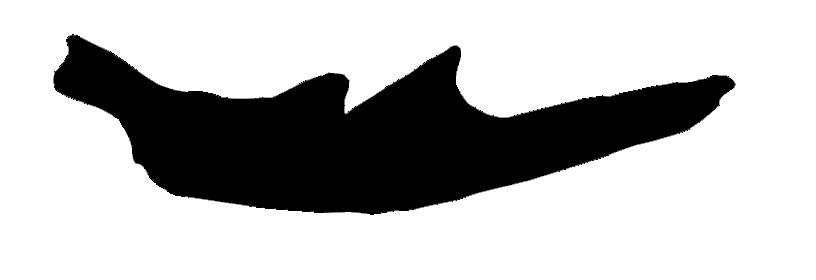

### Att_cuttacutta.jpg

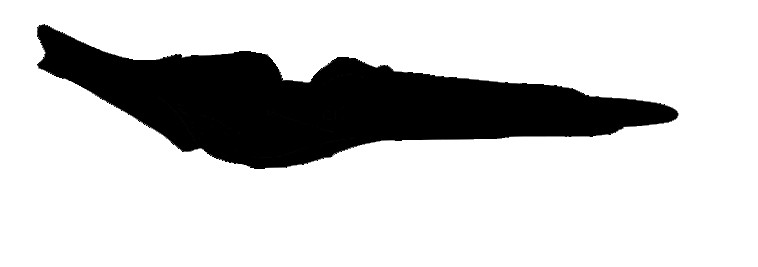

### Att_mainae.jpg

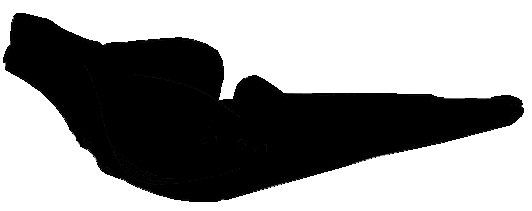

### Att_radon.JPG

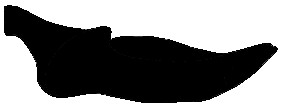

### Baa_firstmani.jpg

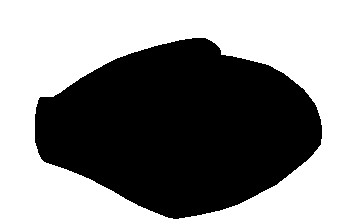

### Baa_magico.jpg

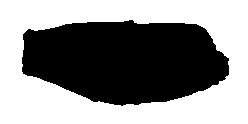

### Bam_bamaga.jpeg

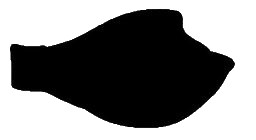

### Bam_milloti.jpg

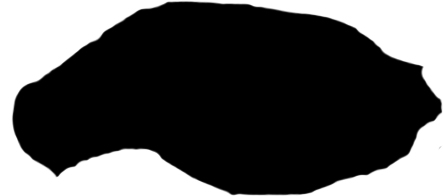

### Bam_pileti.jpg

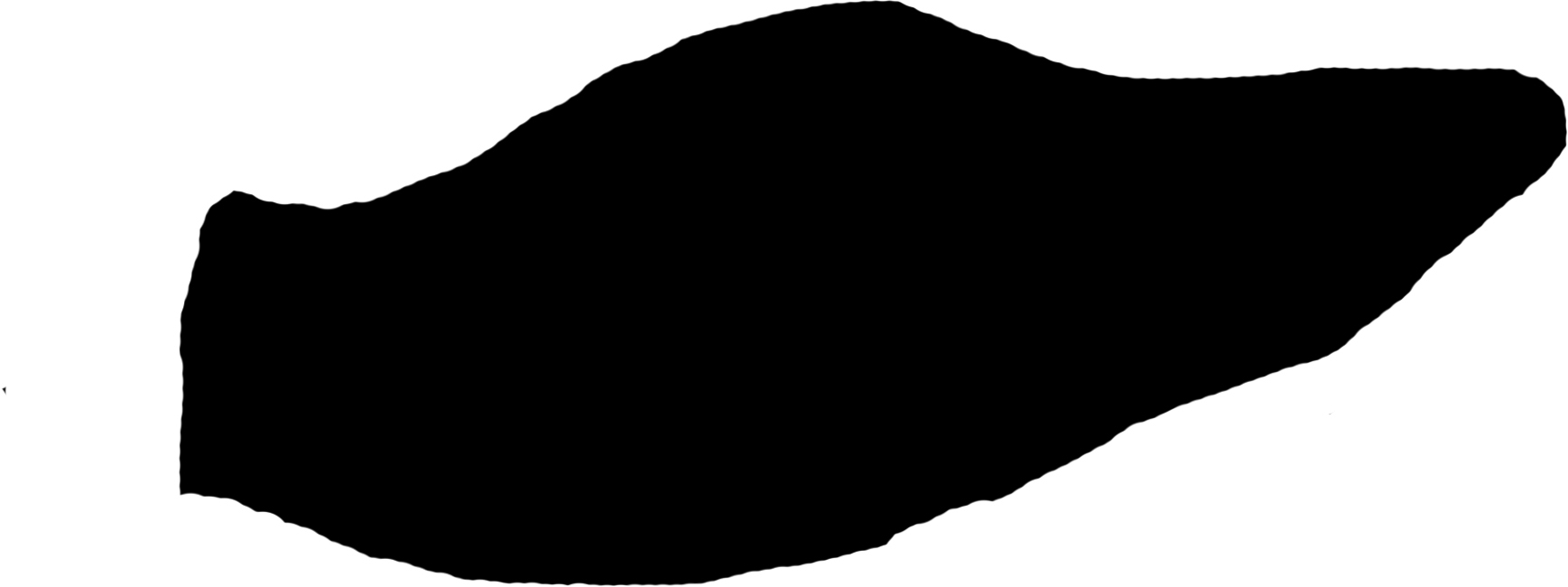

### Bam_siamensis.jpg

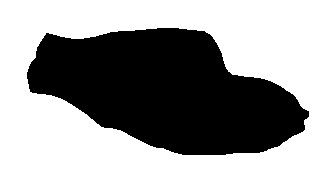

### Bam_subsolanus.jpeg

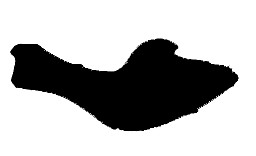

### Bam_vespertinus.jpeg

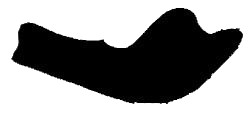

### Bam_weipa.jpeg

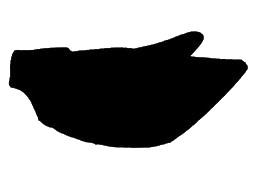

### Bel_pentalatus.jpg

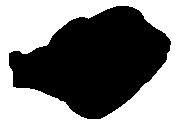

### Bri_nob.jpeg

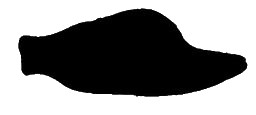

### Bri_walteri.jpg

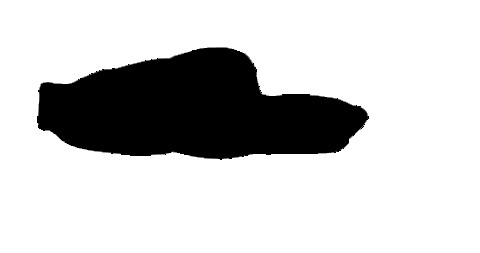

### Bri_woodwardi.jpeg

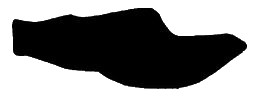

### Buc_hortuspalmarum.jpg

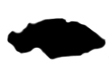

### Cal_bremensis.jpg

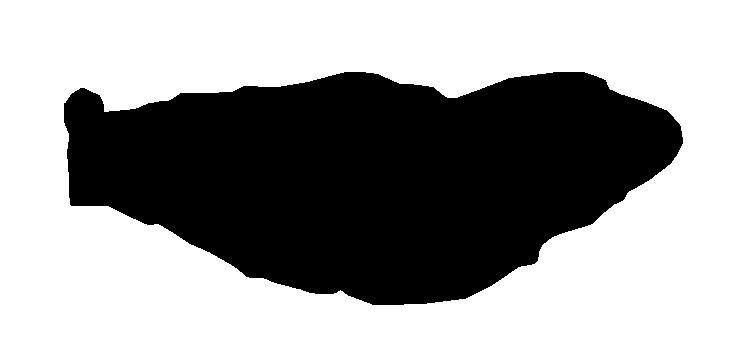

### Cal_embera.jpg

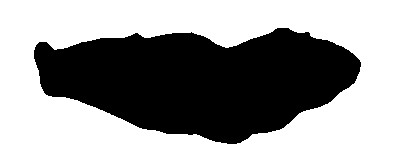

### Cal_nutabe.jpg

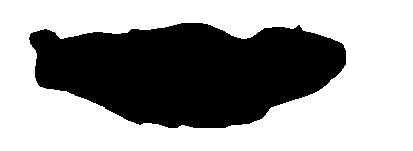

### Cal_valenciorum.JPG

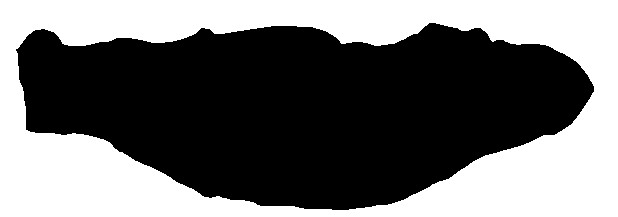

### Can_xikrin.jpg

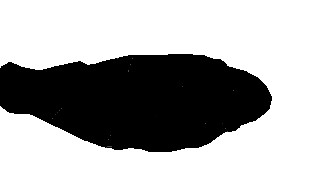

### Car_laurae.jpg

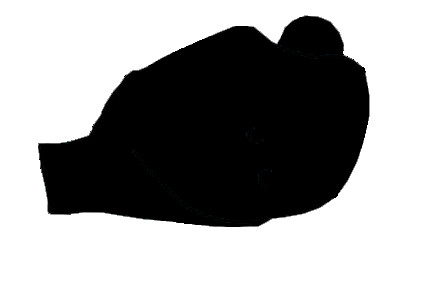

### Cla_claviger.jpg

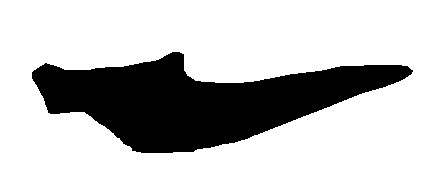

### Cok_ramosiTer07.jpg

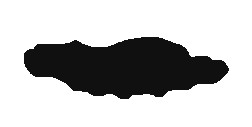

### Col_quindio.jpg

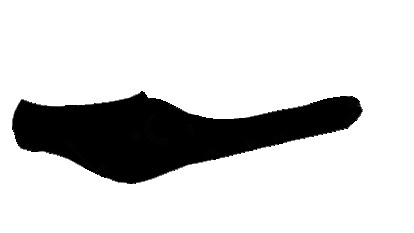

### Col_truncatus.jpg

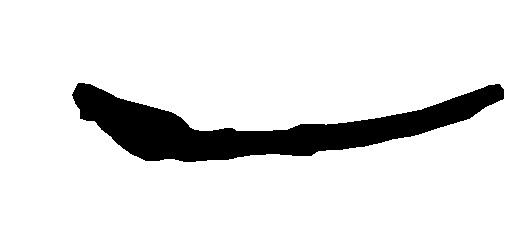

### Cut_rowlandi.jpg

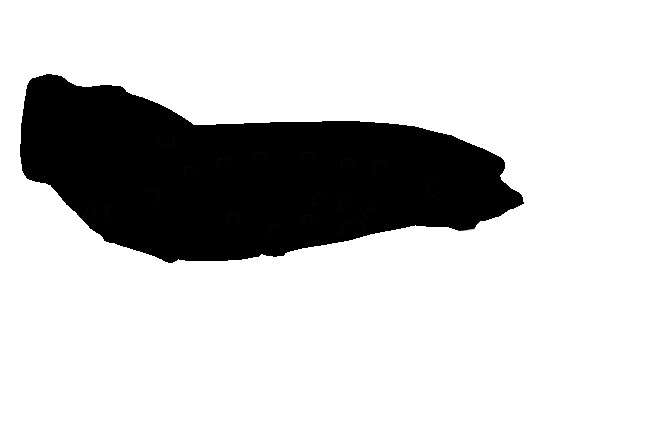

### Cuz_armasi.jpg

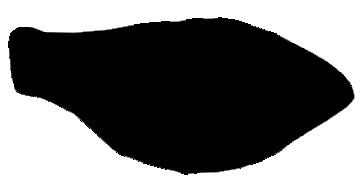

### Cuz_montanusTer04.jpg

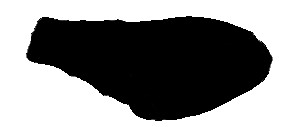

### Cuz_montanusTer17.jpg

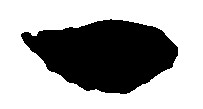

### Cuz_sheylae.jpg

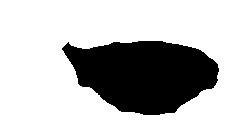

### Dra_anachoretus.jpeg

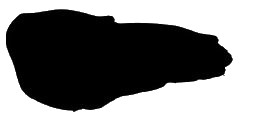

### Dra_belalugosii.jpeg

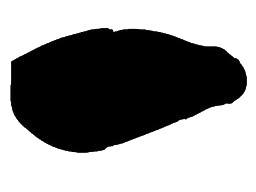

### Dra_bramstokeri.jpg

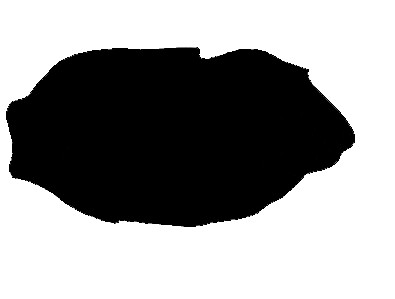

### Dra_brooksi.jpg

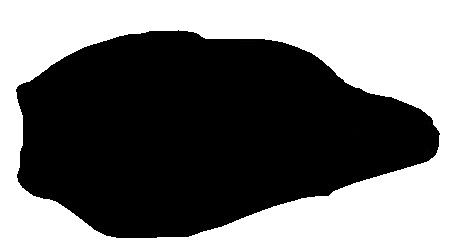

### Dra_bythius.jpg

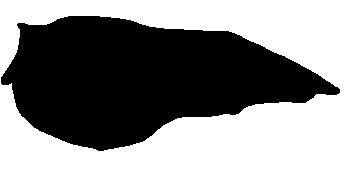

### Dra_christopherleei.JPG

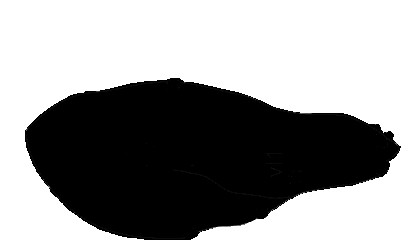

### Dra_eremius.jpg

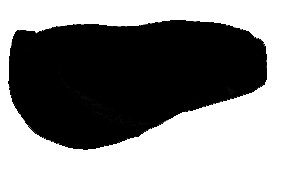

### Dra_gnophicola.JPG

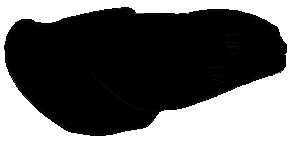

### Dra_julianneae.jpg

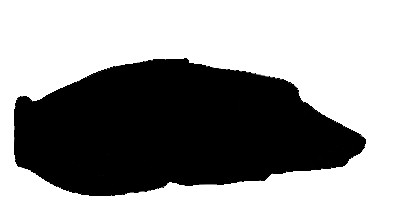

### Dra_karenbassettae.jpeg

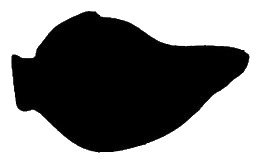

### Dra_kryptus.jpg

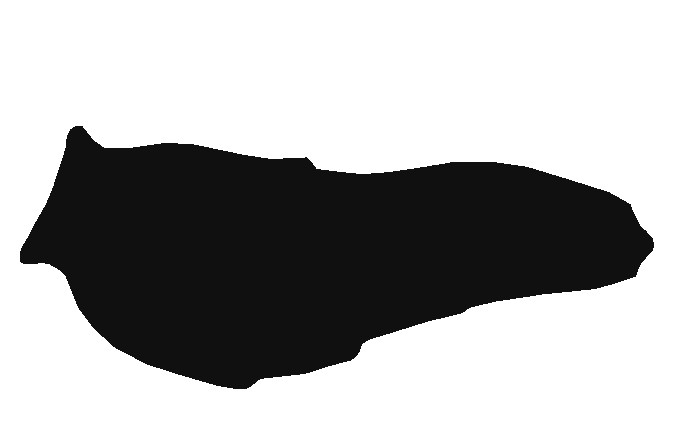

### Dra_mckechnieorum.JPG

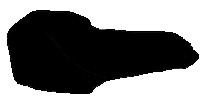

### Dra_neoanthropus.jpg

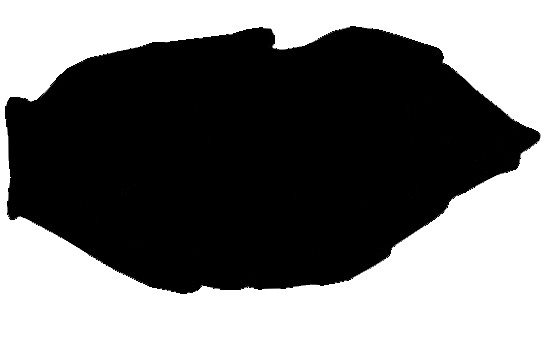

### Dra_vinei.jpg

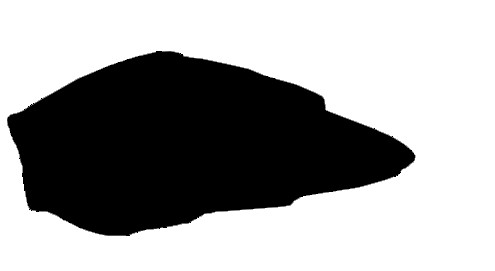

### Dra_warramboo.JPG

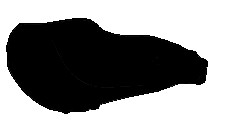

### Dum_decuiRR79.jpg

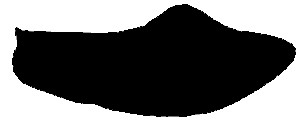

### Eni_eruptoclausus.jpg

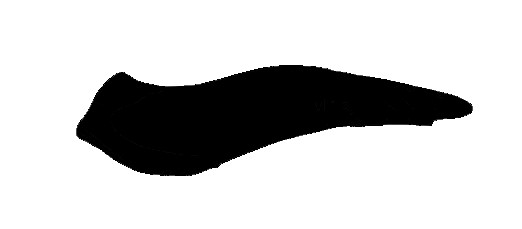

### Gen_sp1.jpg

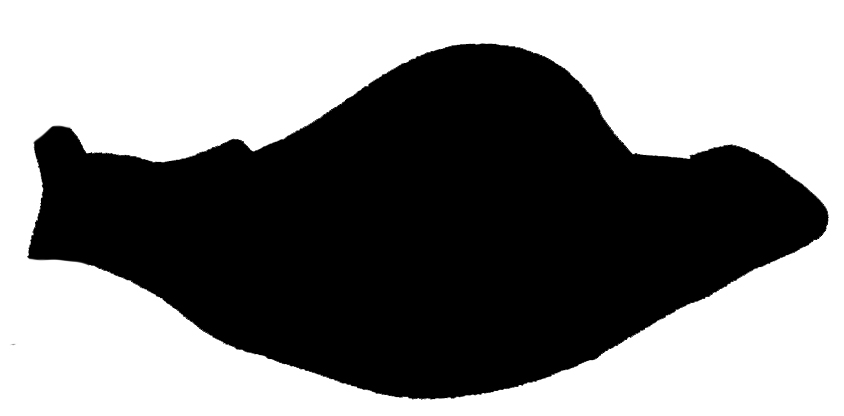

### Gen_sp2.jpg

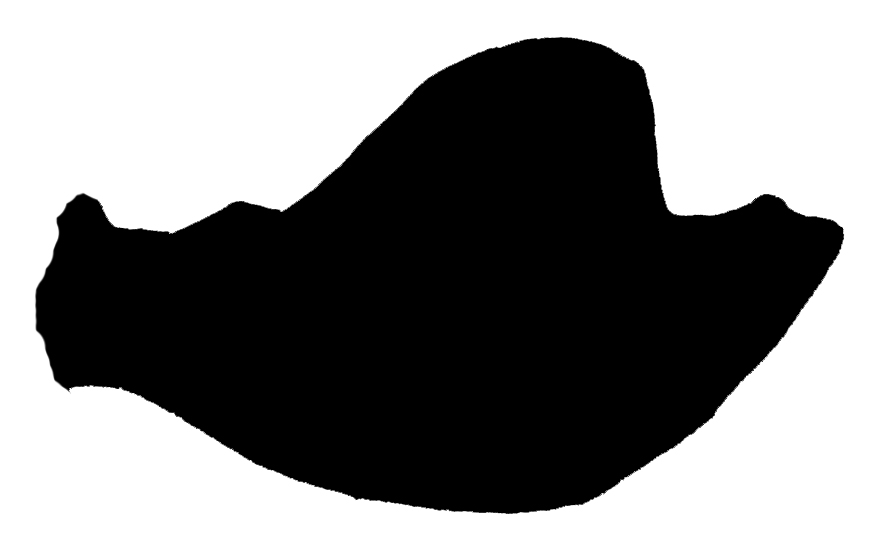

### Gen_sp3.jpg

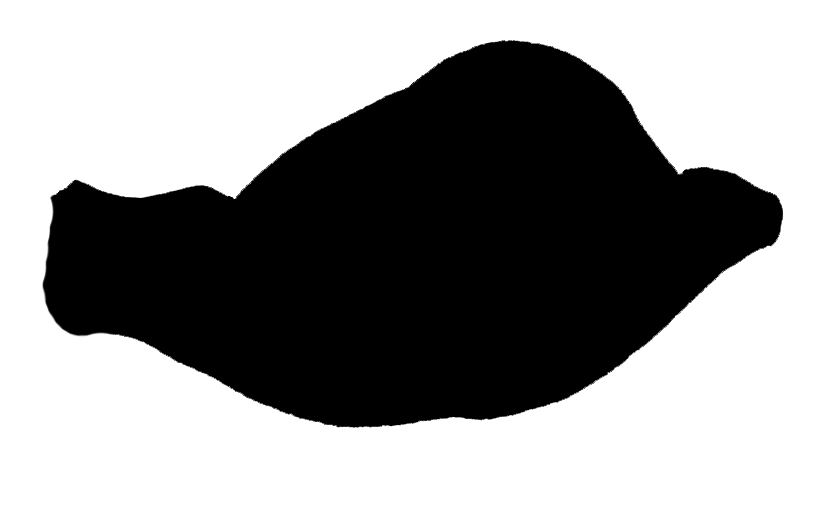

### Gua_armatus.jpg

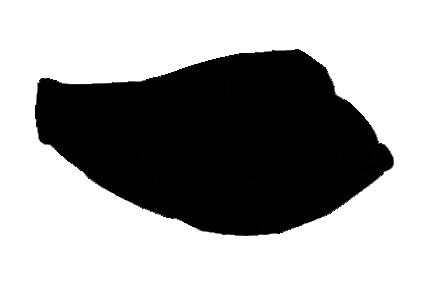

### Han_acrocaudatus.jpg

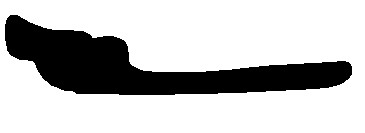

### Han_centralis.jpg

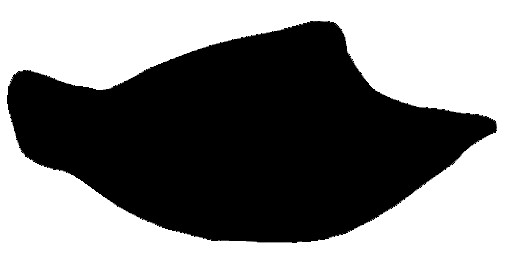

### Han_humbertoi.jpg

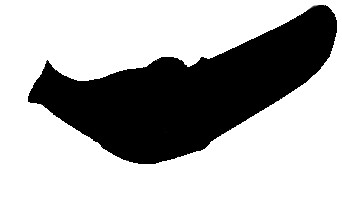

### Han_mumai.jpg

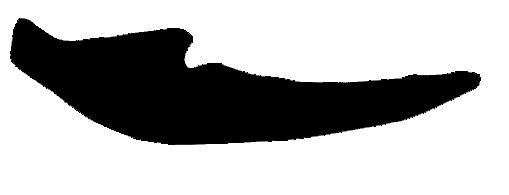

### Han_selva.jpg

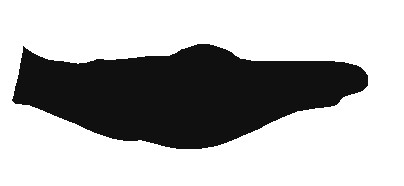

### Han_simonisHS05.jpg

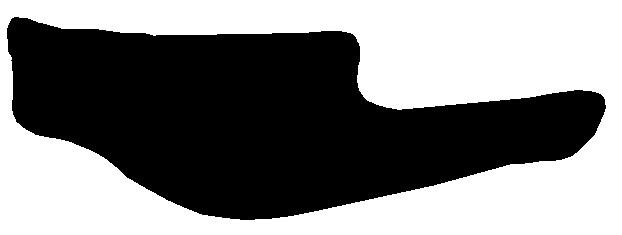

### Han_simonisRR79.jpg

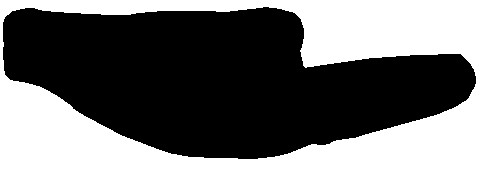

### Han_tobago.jpg

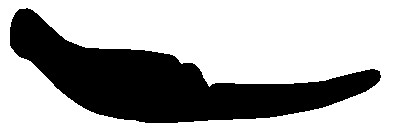

### Han_trinidanus.jpg

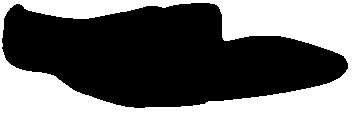

### Han_urbanii.jpg

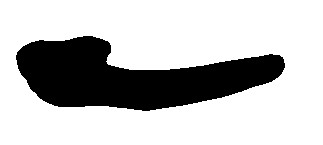

### Han_yolandae.jpg

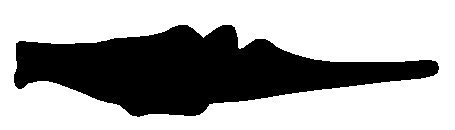

### Row_insignis_H&S_05_dorsal_m.jpg

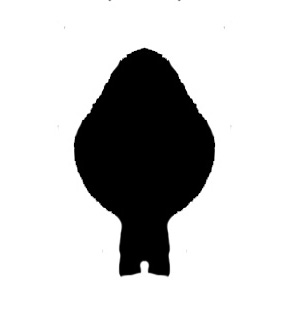

### Row_isabel_Armas_Antunn_02_dorsal_m.jpg

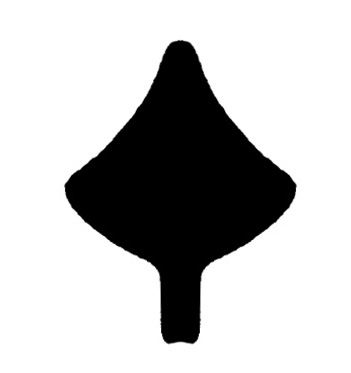

### Row_jarmillae_deAC01_dorsal_m.jpg

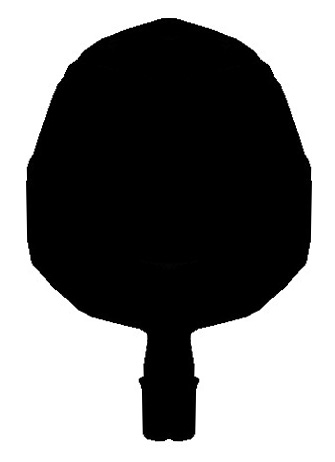

### Row_lantiguai_deArmas04_dorsal_m.jpg

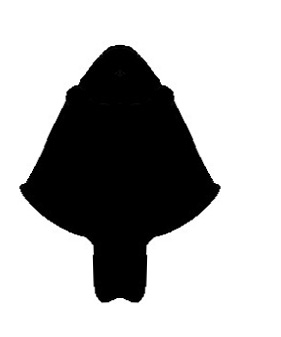

### Row_linsduarteae_S08_dorsal_m.jpg

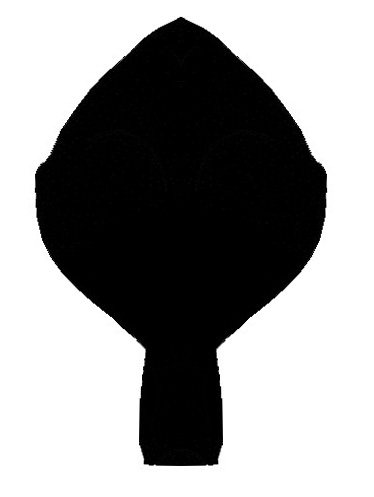

### Row_littoralis_Ter03_dorsal_m.jpg

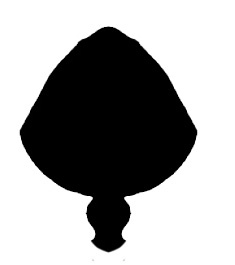

### Row_longipalpus_R&R79_dorsal_m.jpg

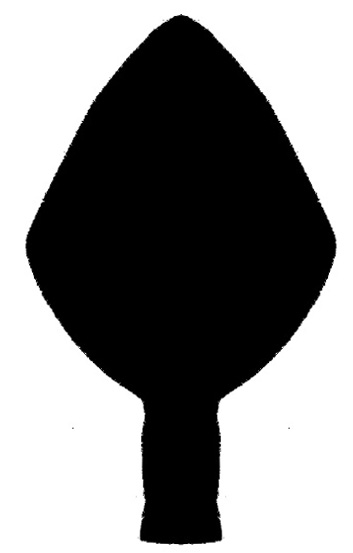

### Row_marianae_Ter03_dorsal_m.jpg

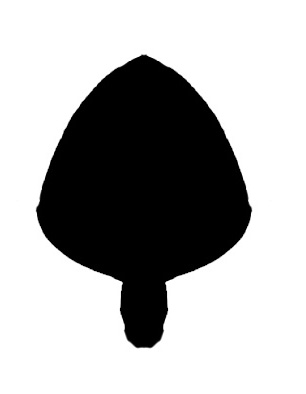
