## Supplementary figures and images for "Tales of schizomid tails: patterns in schizomid flagellum shape from elliptical Fourier analysis"

### Row_longipalpus.jpg

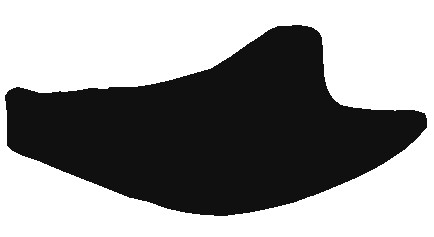

### Row_marianae.jpg

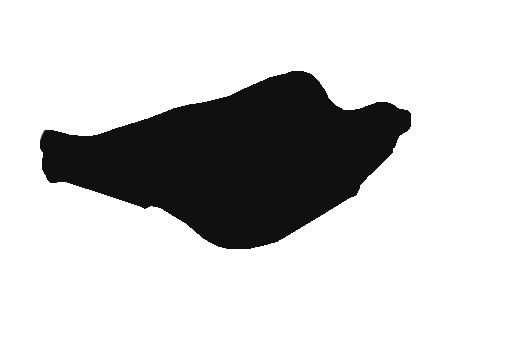

### Row_martinezi.jpg

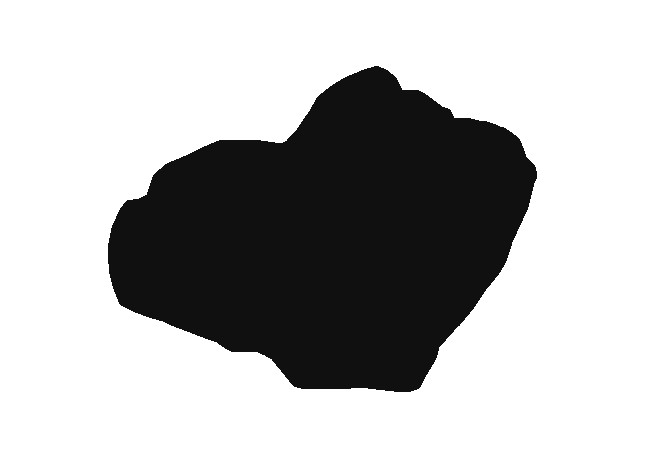

### Row_melici.jpg

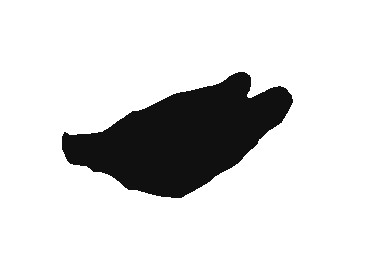

### Row_mixtus.jpg

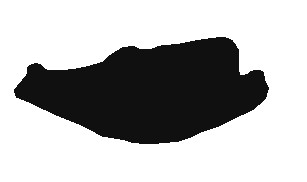

### Row_moa.jpg

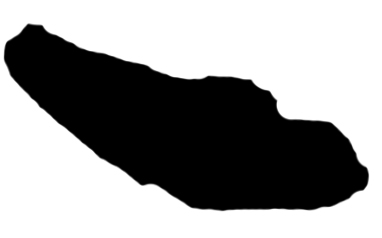

### Row_monensis.jpg

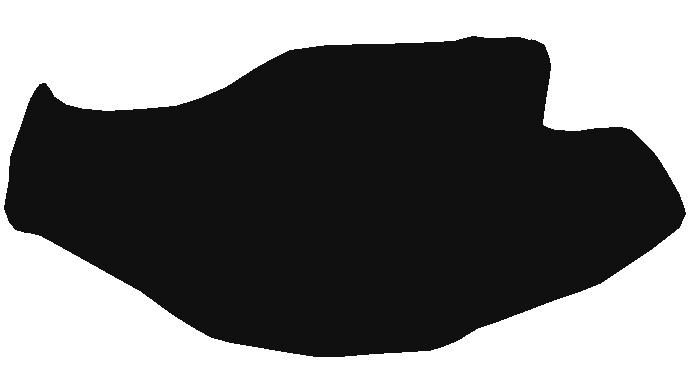

### Row_monticola.jpg

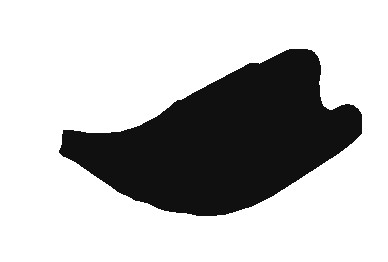

### Row_naranjo.jpg

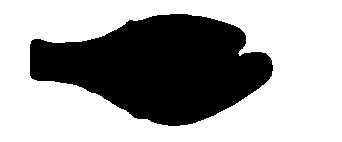

### Row_peckorum.jpg

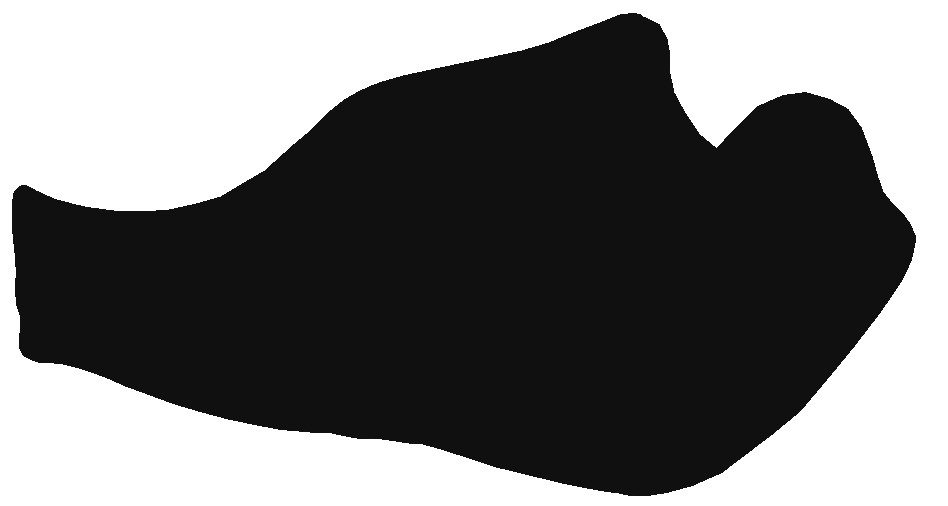

### Row_pedrosoi.jpg

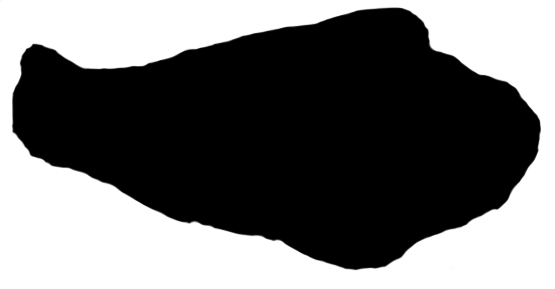

### Row_potiguar.jpg

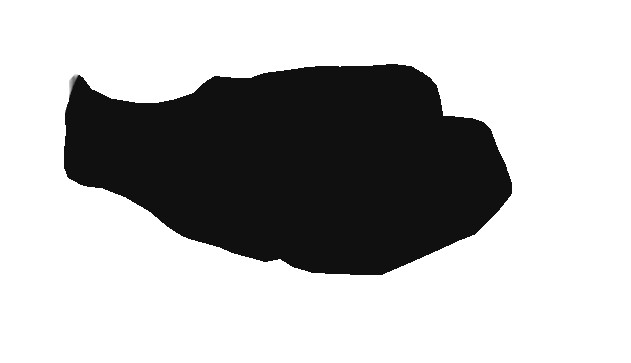

### Supplemental File 11

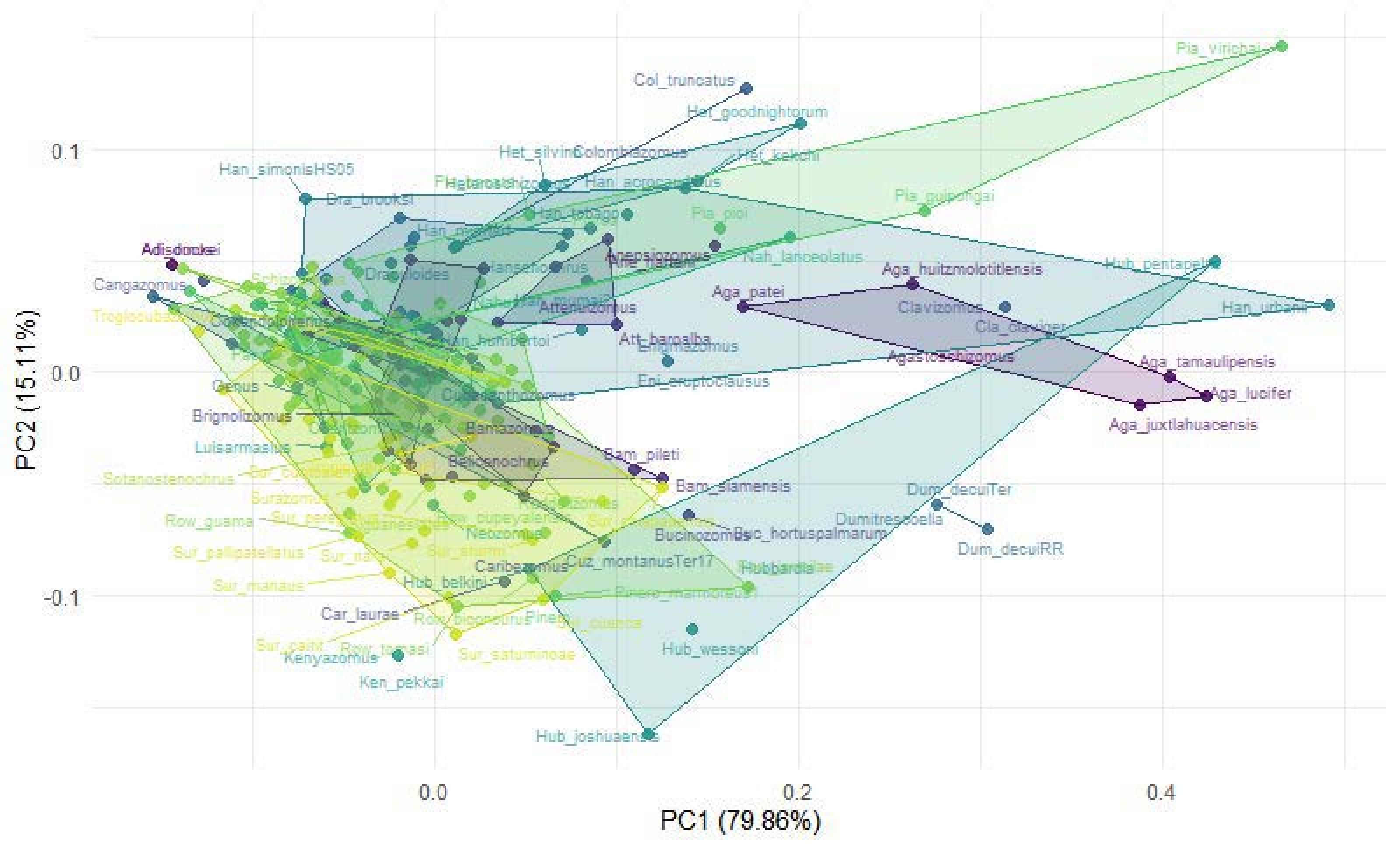

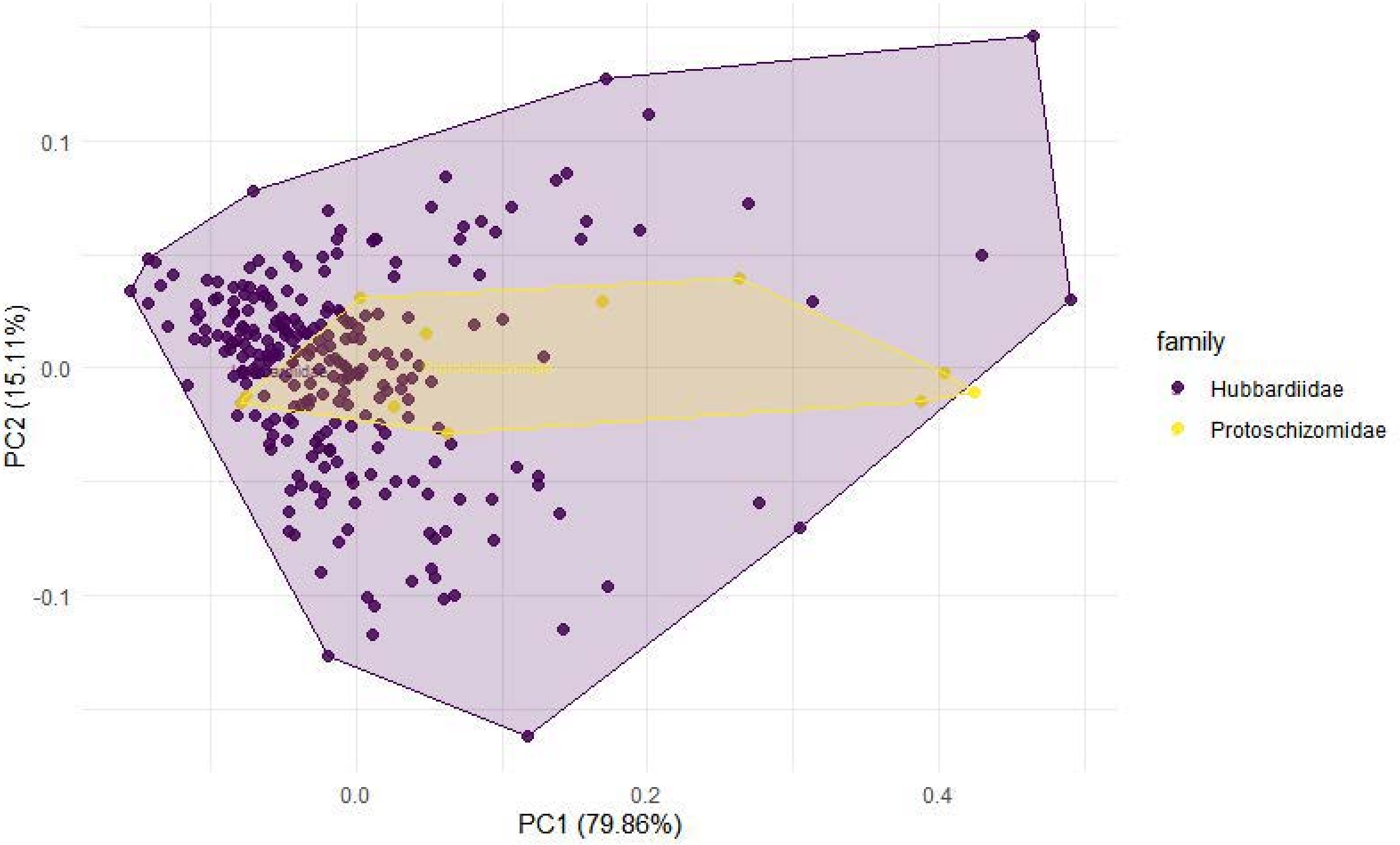

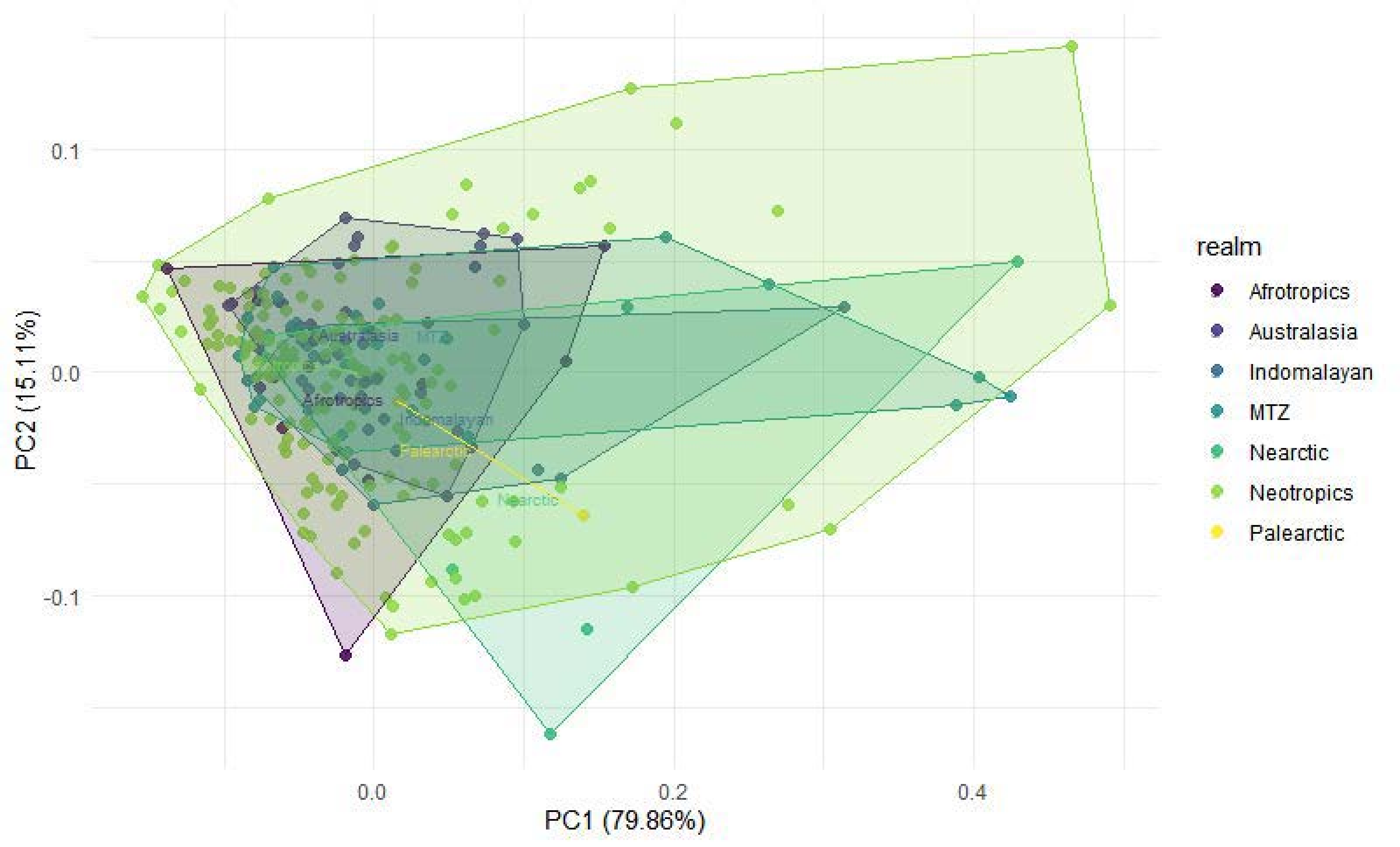

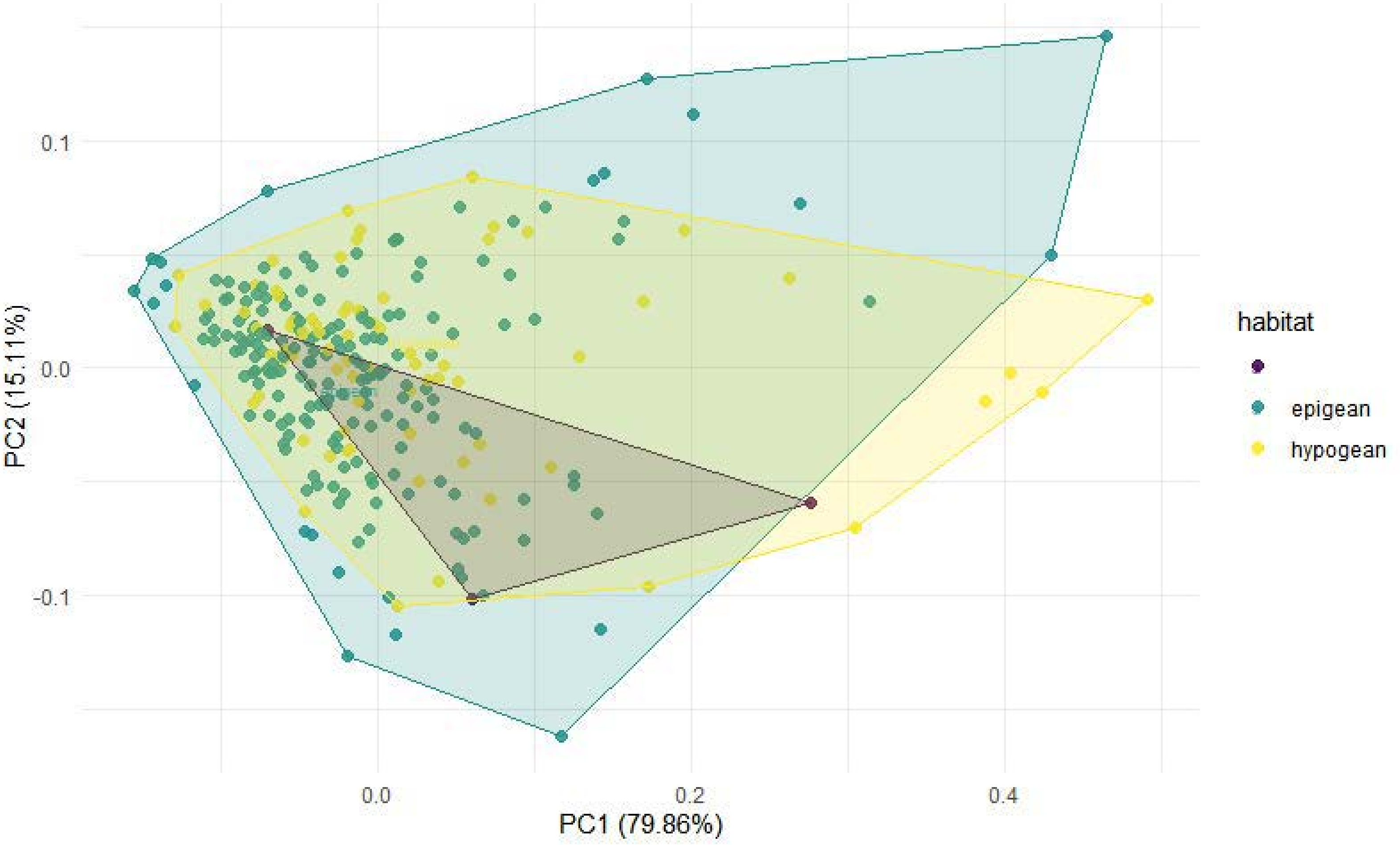

### Supplemental File 12

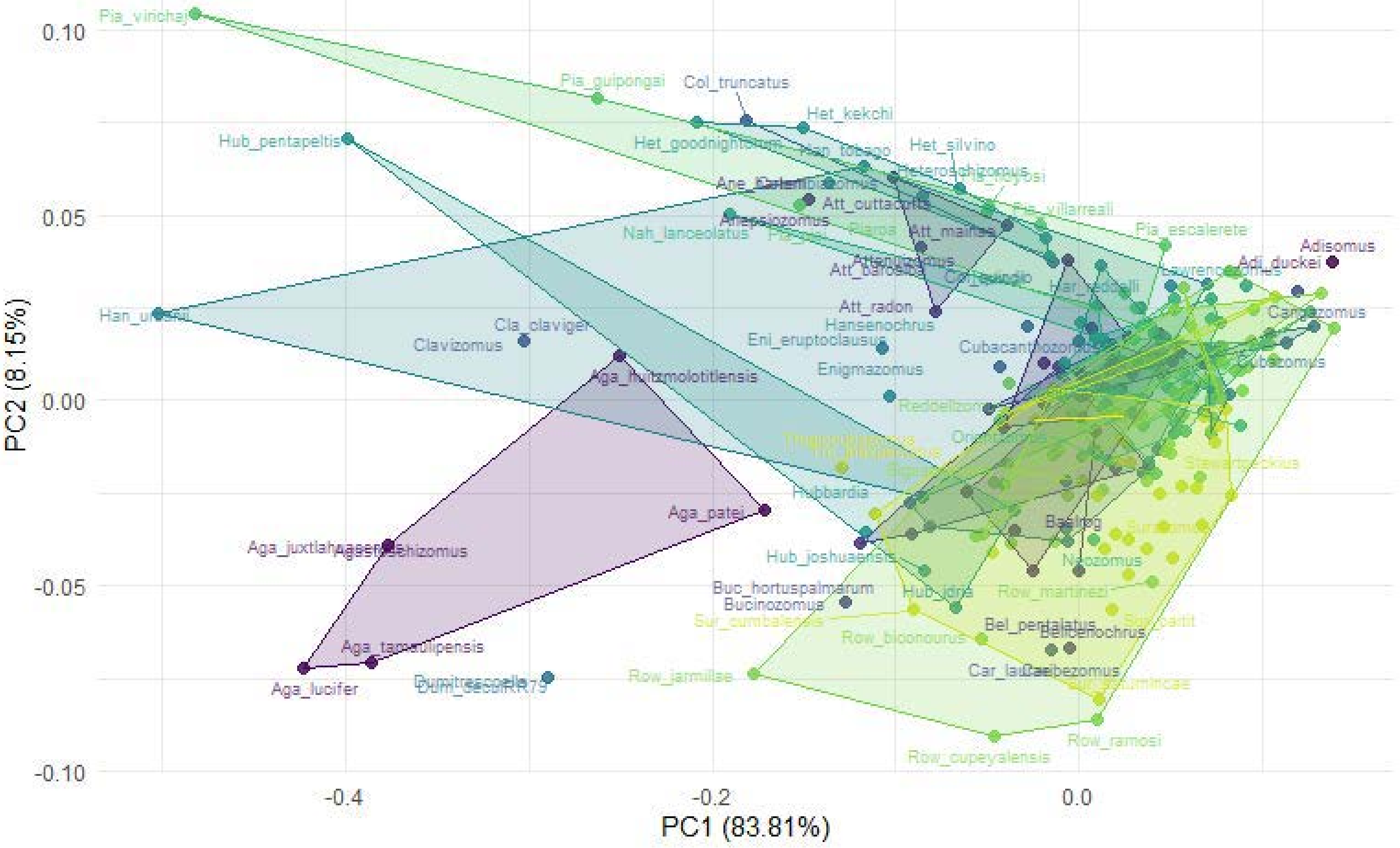

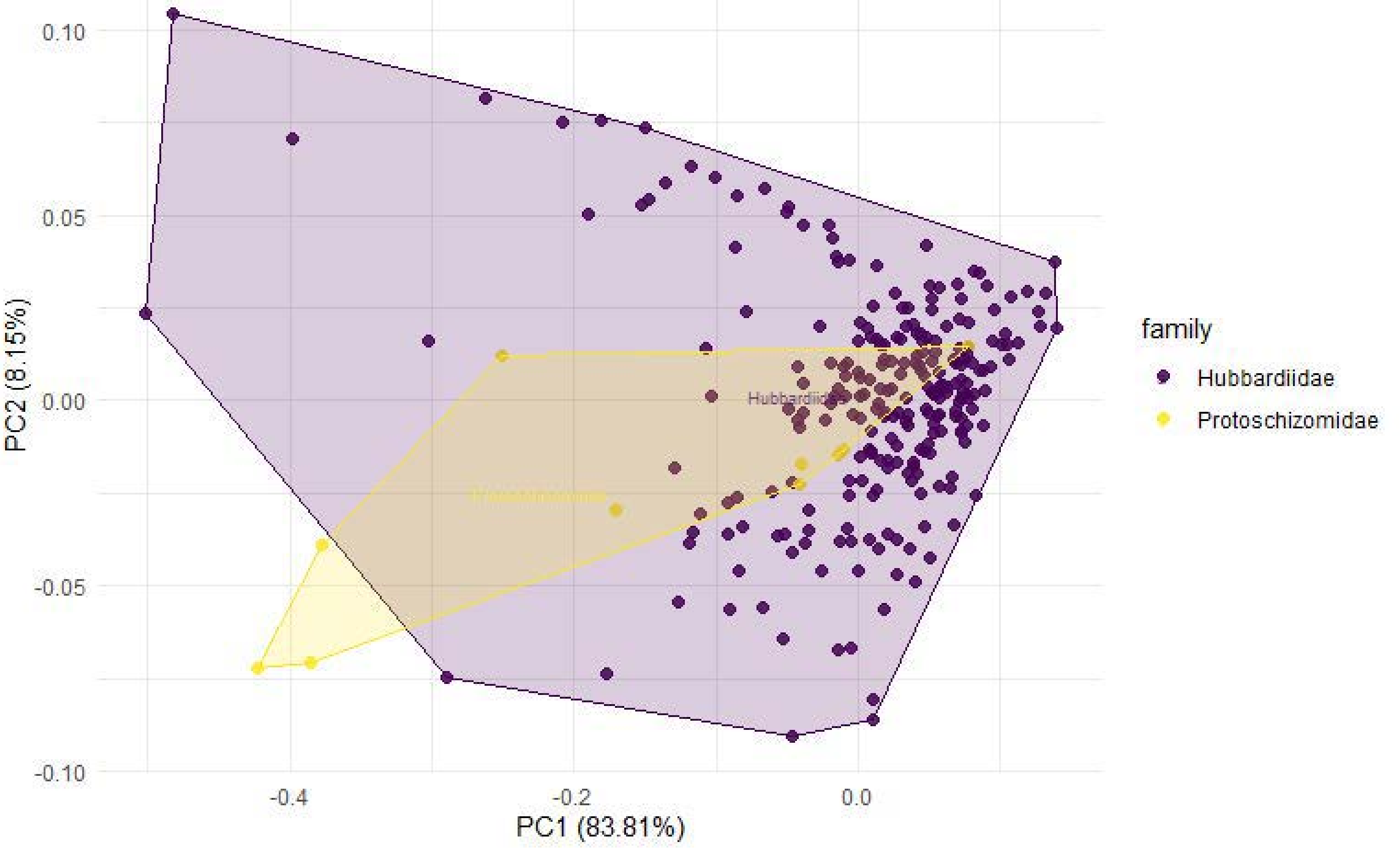

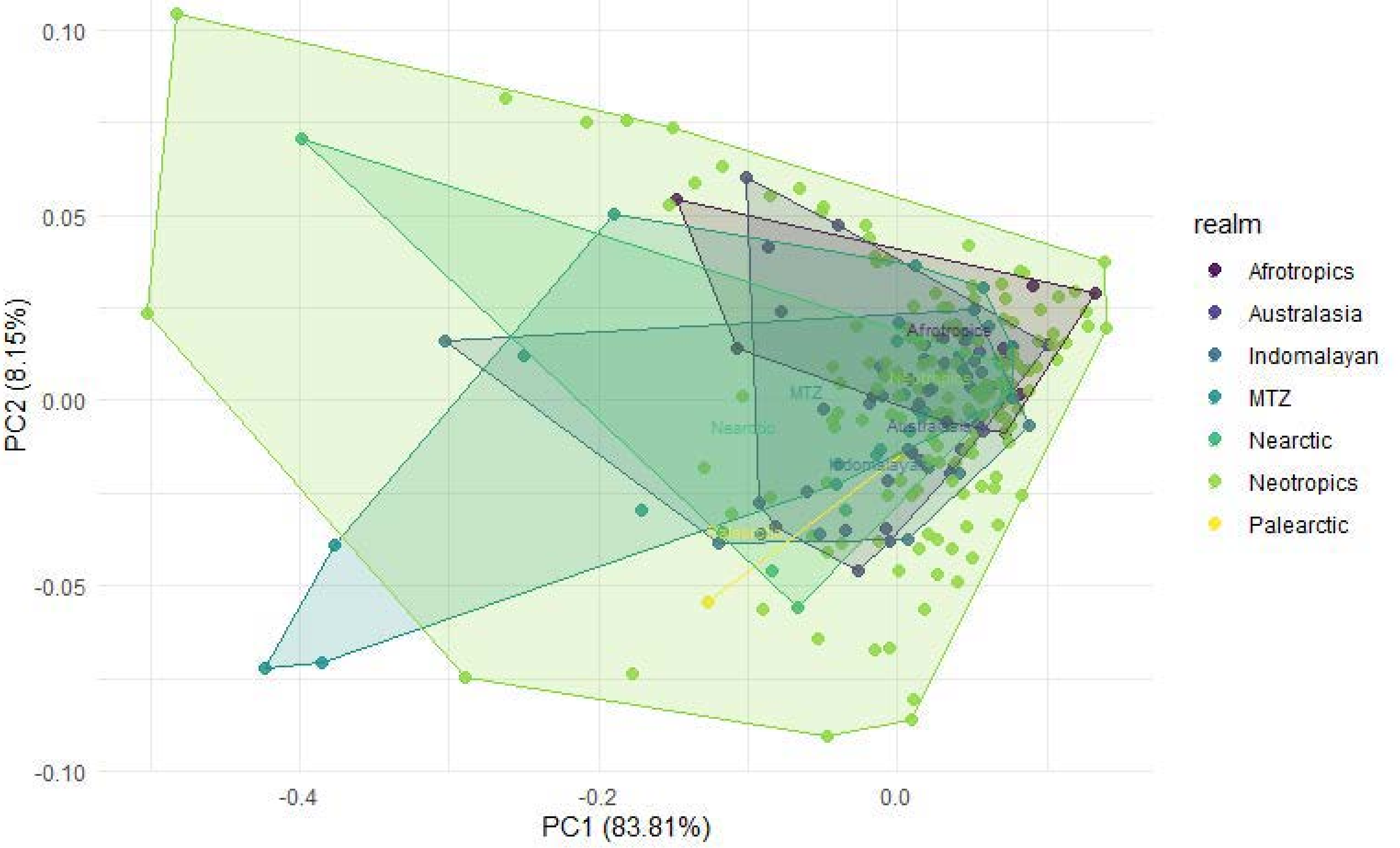

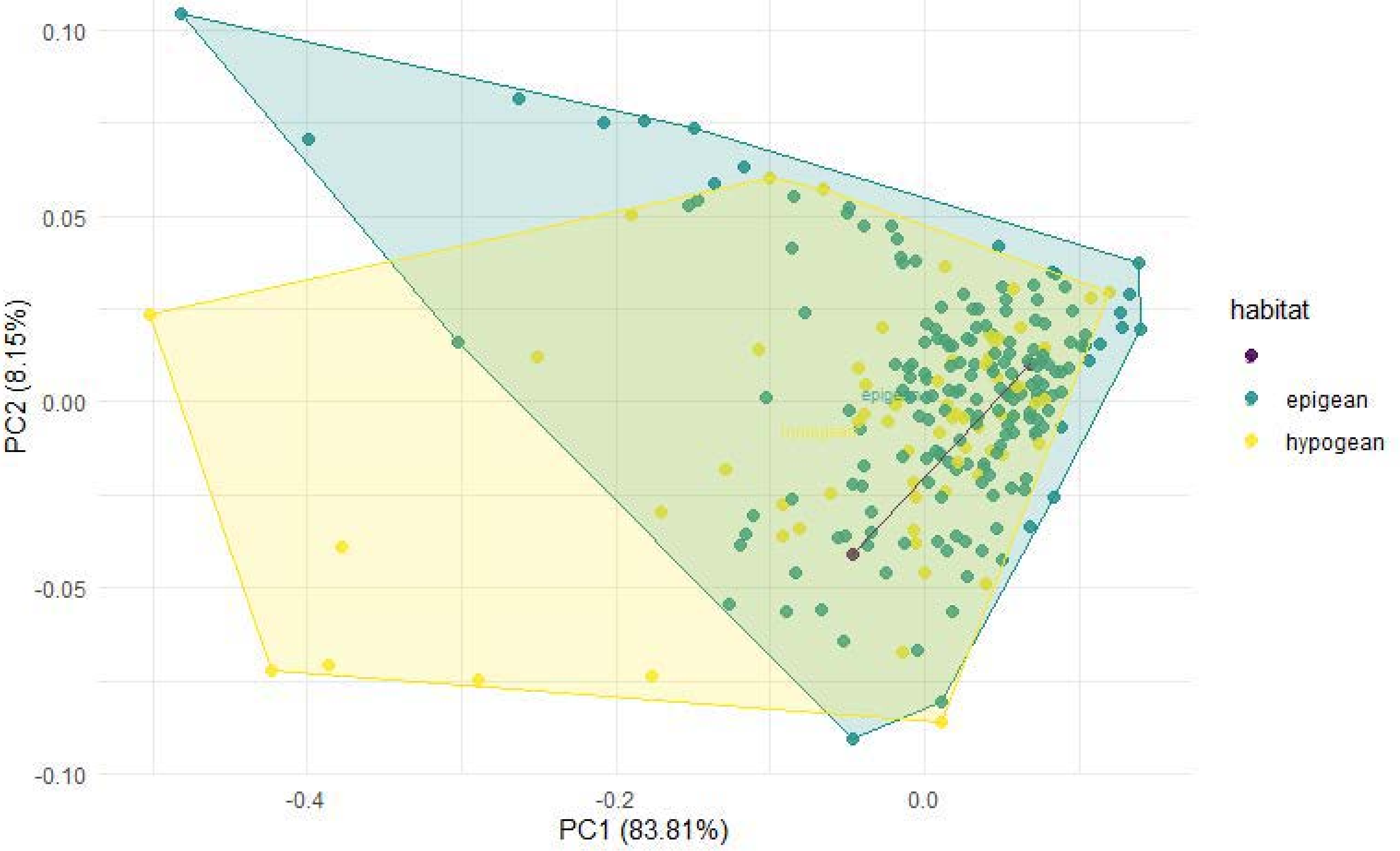

### Supplemental File 14

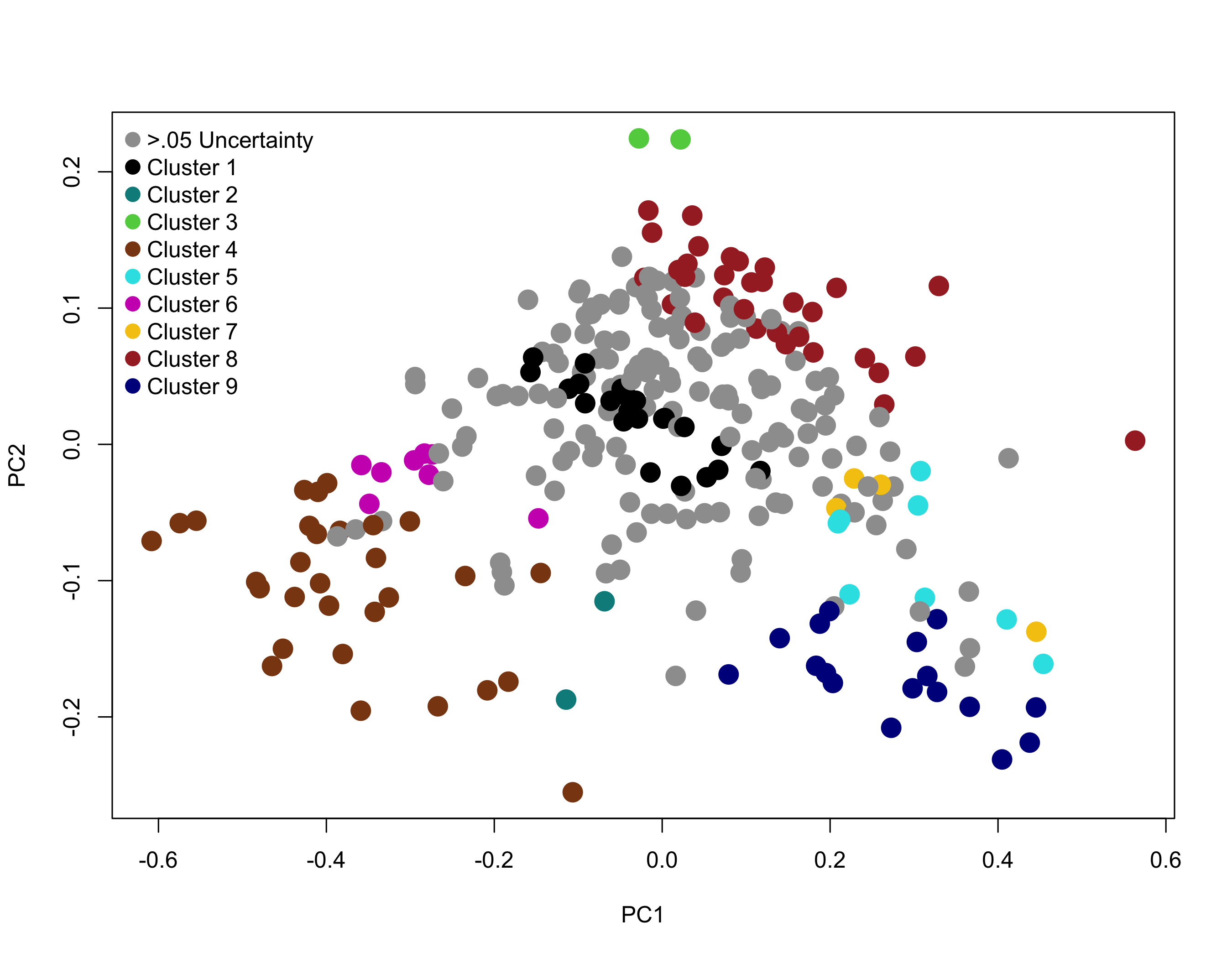
